# Spinal cord regeneration deploys cell-type specific developmental and non-developmental strategies to restore neuron diversity

**DOI:** 10.1101/2025.11.10.687703

**Authors:** Avery Angell Swearer, Samuel B. Perkowski, Iba Husain, Thiago A. Figueiredo, Morgan E. McCartney, Andrea E. Wills

## Abstract

A major goal of spinal cord injury research is to develop a path to endogenous regeneration. This approach has been heavily informed by animal models of natural regeneration. An unresolved question is whether these models rebuild the spinal cord by exclusively accessing developmental mechanisms of neuron differentiation. To address this question, we contrasted single-cell gene expression during regeneration with stage-matched controls in the conditionally regenerative frog *Xenopus tropicalis*. We generated an expanded atlas of neuronal diversity, annotating several neurons in *Xenopus* for the first time. From this atlas, we found that the neuron composition of the developing and regenerating spinal cord differ. So do the strategies employed, which favor waves of cell-type specific neuron morphogenesis, proliferation, and proliferative neurogenesis during regeneration. Low levels of early neurogenesis are then compensated by movement of post-mitotic neurons. Our work highlights the use of distinct developmental versus regenerative paths to heal post-injury.

## Introduction

Spinal cord injury can cause a permanent and devastating impact to quality of life.^1^ Naturally occurring models of vertebrate spinal cord regeneration represent the gold standard for understanding how functionality can be restored after major injury. Such models include zebrafish,^2^ urodeles,^3^ *Xenopus* frogs,^4^ lizards,^5^ and more recently, the spiny mouse *Acomys*.^6^ These studies in animal models are complemented by bioengineering efforts that leverage implanted devices or substrates to encourage neural plasticity or neurogenesis, and by stem cell therapeutic efforts focused on either promoting endogenous repair or exogenous transplantation of neural cell types.^7–9^ All these approaches are heavily informed by our understanding of neurogenesis as it occurs during embryogenesis, where the mechanisms that give rise to spatially distinct neuron domains have been elucidated in beautiful molecular detail.

During embryogenesis, the vertebrate spinal cord is elegantly patterned into distinct neuronal domains from regionally restricted neural progenitor cell (NPC) populations.^10^ Neurogenesis progresses through development in waves, dependent on windows of progenitor specification and differentiation. The spinal cord is thus made up of neurons generated either through primary neurogenesis during neurula stages or ongoing secondary neurogenesis.^11,12^ The early spinal cord includes major neuron types such as dorsal primary Rohon-Beard sensory neurons (RBNs)^13^, various primary and secondary interneurons (INs), ventrolateral motor neurons (MNs), and ventral secondary Kolmer-Agduhr interneurons (KAs, or cerebrospinal fluid-contacting neurons (CSF-cNs) in mammals)^14,15^ (reviewed in *Xenopus* in ^16–18^). In addition, specific cardinal IN types (dI1-6, V0-3) have been documented in zebrafish and mice but have largely not been characterized in *Xenopus* frogs. Overall, the presence and positioning of these neuron types is crucial to forming a complete neural circuit that allows larvae to respond to sensory stimuli via motor movement. The transcriptional regulation of NPC organization and differentiation into INs, KAs, and MNs is highly conserved across vertebrates, although RBNs are only found in fish and larval amphibians and degenerate during later development to make way for the dorsal root ganglia.^19^ Despite the plethora of research describing these neurons during development and homeostasis, little is known about how—and if—each of these neuron types regenerate.

Studies of spinal cord regeneration across animal models have made clear two considerations about development and regeneration. First is that the path to regeneration varies by species, both with respect to the completeness of spinal cord regeneration and with respect to how developmental signals are redeployed. An excellent example is the role of Sonic hedgehog (Shh) signaling, which is an indispensable regulator of dorsal versus ventral cell fate in the developing spinal cord.^20^ While anole lizards fail to regenerate dorsal NPC domains due to ubiquitous shh expression,^5^ axolotls^21,22^ and *Xenopus tropicalis*^23^ reactivate Shh signaling in ventral tissues to recapitulate dorsal/ventral patterning post-injury, although *Xenopus* expresses Shh in the ventral floor plate and notochord and axolotls express shh only in the floor plate in a divergence from developmental patterning. Thus the degree of functional regeneration and extent of developmental recapitulation vary by species, even among regeneratively competent models. The second consideration is that spinal cord regeneration can employ neurogenic mechanisms that differ from developmental neurogenesis. Examples include dedifferentiation of radial glia into neural progenitors, which has been well documented in zebrafish and axolotls,^24,25^ movement of differentiated neurons,^26^ and generation of regeneration-specific, injury-induced neurons (iNeurons) in zebrafish.^27^ These studies have made it evident that many processes can contribute to neural regeneration, but the picture of how these mechanisms underlie the repopulation of diverse neuron types is incomplete.

Most animal studies leverage adult spinal cord transection as their injury model, leaving a gap in our understanding of how complete regeneration of a spinal cord proliferative compares to development. *Xenopus* frogs are a unique model in that they are regenerative during larval stages but lose the ability to undergo spinal cord regeneration during metamorphosis,^28^ suggesting that their regenerative competence at tadpole stages is tied to their access to ongoing developmental mechanisms. *Xenopus tropicalis* is therefore an ideal model in which to compare developmental and regenerative neuron repopulation. Here we used scRNA-seq to establish a molecular atlas of neuron types in the tadpole, then defined how the neuronal cell composition of the spinal cord changes during early, intermediate, and late phases of regeneration. We specifically contrasted the cell composition and organization of the post-injury spinal cord as regeneration progresses with that of the stage-matched developing spinal cord. We then delved more deeply into how the many neurogenic mechanisms that have been described in regeneration are equipped by distinct neuronal cell types, using HCR and lineage tracing to elucidate distinct roles of proliferative neurogenesis and post-mitotic neuron movement.

## Results

### scRNA-seq shows dynamic changes in cell type abundance over developmental and regenerative time

Our principal goal was to catalog changes in neural cell type composition in the spinal cord during tail regeneration and compare these with changes over a corresponding developmental timeline. To this end, we performed single-cell RNA sequencing on injured and stage-matched uninjured tail tissue from 0 to 7 days post amputation (dpa), beginning at stage 41 (3 days post fertilization).^29^ By 7 dpa, tail length in injured tadpoles matched uninjured peers and we considered regeneration complete (Figure S1A, B). To enrich for neural cells we used the transgenic line Xtr.Tg(*pax6*:GFP;cryga:RFP;*actc1*:RFP), which broadly marks NPCs with GFP in *X. tropicalis*.^30^ At stage 41, we amputated tadpoles and collected the posterior third of the tail as our “tail tip” sample and ∼100 microns of tissue proximal to the amputation plane as our “mid-trunk” sample to control for anterior/posterior differences in cell composition (Figure 1A). We then split tadpoles into uninjured and regenerating groups and collected tail tissue from both groups at 1 dpa, 3 dpa, and 7 dpa to represent early, intermediate, and late spinal cord regeneration, respectively (Figure 1A). Permissive FACS gating conditions were used to enrich for high-expressing *pax6*:GFP+ cells (such as NPCs) as well as low-expressing *pax6*:GFP+ (such as differentiated neurons) and some *pax6*:GFP-cells, allowing us to populate non-neural and non-*pax6* clusters. After quality control filtering, a total of 20,450 uninjured and 15,402 regenerating cells were included for downstream analysis with Seurat (Figure S1C).^31–35^ After dataset integration and UMAP dimensional reduction we identified clusters, to which we assigned unique cell type labels based on previous scRNA-seq analyses in *Xenopus* and other relevant developmental markers (Figure 1B, Figure S2).^36,37^

**Figure 1.**
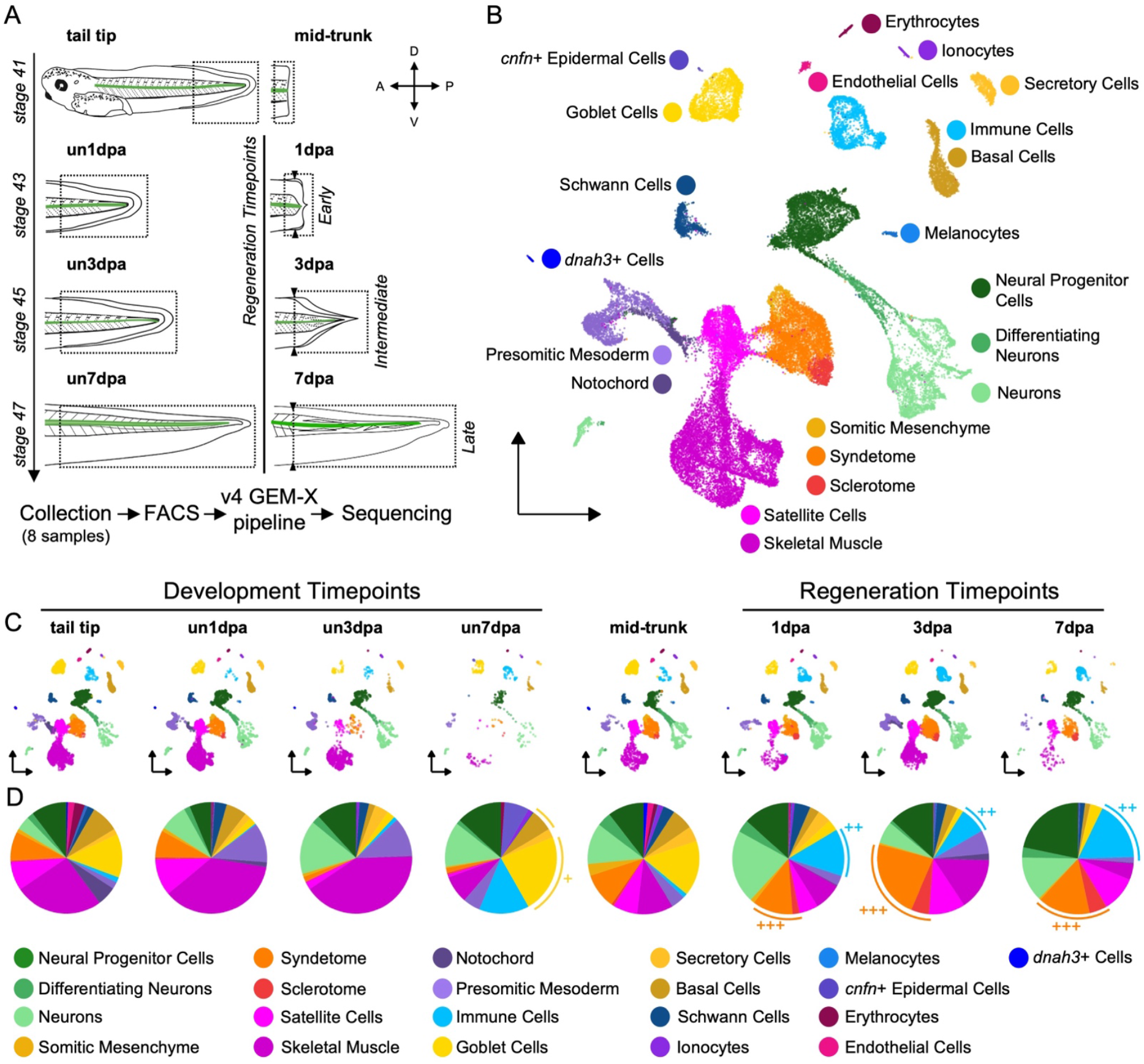
Single-cell RNA sequencing reveals dynamic changes in non-neural cell populations during development and regeneration. A) Experimental design of single-cell RNA sequencing experiment, starting with the tail tip (stage 41, posterior third of the tail) and mid-trunk (100 µm proximal tissue to amputation plane) before following up with developmental uninjured (un) or amputated samples at 1, 3, or 7 days post amputation (dpa). Samples were collected and sorted via FACS before processing and sequencing. B) UMAP of 35,852 cells grouped into 21 neural and non-neural cell clusters. C) UMAP of each timepoint, including developmental and regenerating samples. D) Pie chart showing the cluster makeup of each sample. ^+^denotes expanded goblet cells and basal cells in development, ^++^denotes expanded immune cells in regeneration, and ^+++^denotes expanded syndetome and sclerotome in regeneration.

We first examined global trends in cell type composition as tail development and regeneration progressed. Many of the patterns we observe for days 1-3 are similar to those observed in *X. laevis*.^36^ While the overall cell type composition of tail tips and mid-trunk tissue was similar, we noted several striking changes in cell type abundance between the stage-matched developmental and regenerating timepoints by 7 dpa (Figure 1C). Examples include the expansion of goblet cells and basal cells in development but not regeneration (Figure 1D^+^), and the relative enrichment of immune cells (Figure 1D^++^) and immature connective tissue types like sclerotome and syndetome (Figure 1D^+++^) in regeneration.

### An expanded atlas of Xenopus neuron types based on conserved marker genes

For the remainder of our study, we focused on changes in neural cell types. To interrogate spinal cord regeneration, we subset 7,337 neural cells (3,708 uninjured cells and 3,629 regenerating cells). We then set out to use molecular markers to better define a molecular atlas of neuron types. Previous designations of larval neuron types in *Xenopus* have relied on their morphology and reactivity. These approaches have enabled the identification of dorsal Rohon Beard Neurons (RBNs),^13^ intermediate excitatory and inhibitory Interneurons (INs), ventrolateral motor neurons (MNs), and ventral cerebrospinal fluid contacting Kolmer-Agduhr interneurons (KAs)^14,15^ (Figure 2A, reviewed in ^16–18^). Three subtypes of interneurons have subsequently been identified based on their expression of *vsx2, engrailed1*, or *evx1*.^14,38,39^ In our analysis, we first mapped clusters back to *Xenopus* neuron types based on known neurotransmitter types and markers, where available (Figure 2B). We then used unique patterns of gene expression from zebrafish and mouse to assign identities to neuron classes that had not previously been molecularly classified (Figure 2A-C, Table S1).

**Figure 2.**
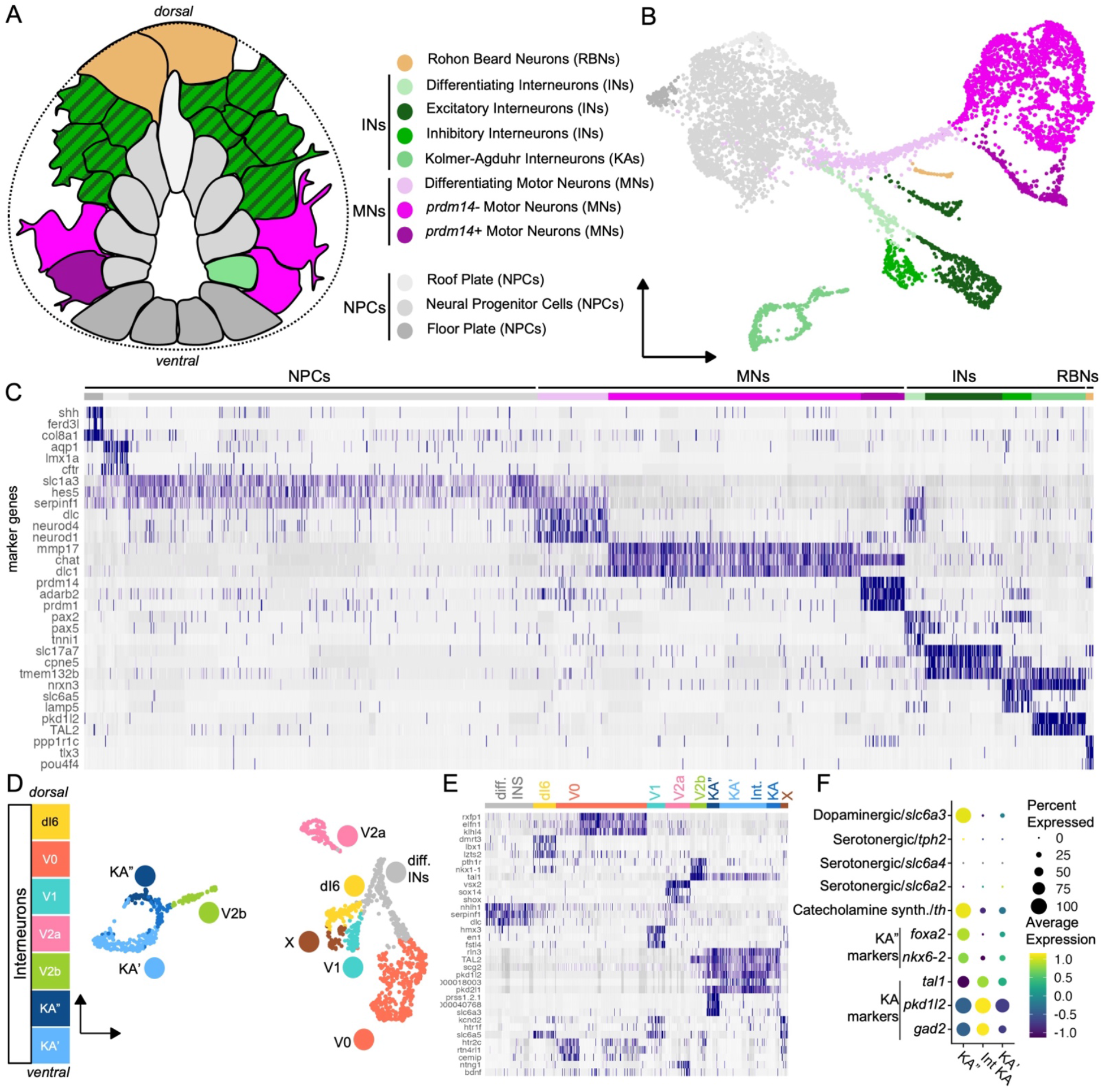
Distinct major and cardinal neurons can be identified using conserved markers in the *Xenopus tropicalis* spinal cord. A) Cross-section diagram showing the relative position of major spinal cord cell types in a stage 41 tadpole. B) UMAP of the merged neural subset showing major NPC and neuron cell types. C) Top 3 markers of each cell cluster. D) UMAP and legend of subset cardinal interneuron types in dorsal/ventral order. In addition to differentiating interneurons (diff. INs) Interneuron populations include KA’ and KA” clusters, a cluster corresponding to V2b inhibitory interneurons (Ventral Longitudinal Descending or VeLD in zebrafish), a cluster corresponding to V2a Excitatory interneurons (Circumferential Descending (CiD) interneurons in zebrafish), a cluster corresponding to V1 inhibitory interneurons (Circumferential Ascending (CiA) interneurons in zebrafish), a cluster corresponding to V0v excitatory interneurons (Multipolar Commissural Descending and Unipolar Commissural Descending or MCoD/UCoD in zebrafish), and a cluster corresponding to inhibitory dI6 interneurons (Commissural Local or CoLo in zebrafish). These types were determined by (E) the top 3 gene markers of each interneuron type and marker genes (Table S1). X denotes unidentifiable glycinergic interneuron groups. F) Dot plot showing gene expression between KA”, intermediate Interneuron (Int KA), and KA’ interneuron groups.

Through this combined approach of validation and conservation, we identified RBNs as well as several types of both known and new MNs, INs, and KAs in our merged dataset. We found two main subtypes of MNs in our dataset: a *prdm14*+, *adarb2*+, *prdm1*+ cluster, and *prdm14*-, *dlc1*+, *mmp17*+ cluster (Figure 2C). *prdm14* is thought to play a role in primary MN differentiation in zebrafish^40^ and may represent differences in the birth time or innervation of these MN clusters. Within the IN cluster, we confirmed the presence of *en1*+ inhibitory GABAergic/glycinergic INs, *vsx2*+ excitatory glutamatergic/cholinergic INs, and *evx1*+ excitatory glutamatergic INs as previously identified in *Xenopus laevis*.^15,38,39^ These populations likely correspond with V1, V2a, and V0v INs, which have been morphologically identified as ascending or descending INs and commissural INs in *Xenopus laevis*, respectively (Figure 2D-E, Table S1). In addition, for the first time in *Xenopus* we found populations of descending INs corresponding to V2b inhibitory INs and dI6 INs that likely represent commissural INs (Figure 2D-E, Table S1). While other cardinal classes such as V3 and dI1-5 could not be identified, it is not yet clear if these neuron types are not present in *Xenopus tropicalis* at these stages or if they arise in different temporal or spatial locations than our collection points. Finally, one cluster was identified as differentiating INs and one cluster of glycinergic INs remained unidentifiable (Figure 2D-E).

Interestingly, the inhibitory KA INs split into 2 mature clusters and one intermediate cluster. Two subtypes of KAs, KA’ and KA’’, have been reported in zebrafish, mice, and other species but not previously in *Xenopus*.^41^ In our dataset, both clusters express *gad2* and *pkd1l2*, but only one expresses KA” markers *foxa2, nkx6-2, and th (tyrosine hydroxylase)*. Given that the *th* expressing cluster did not express other markers of catecholaminergic or serotonergic activity (*slc6a2, slc6a4, tph2*) and did express *slc6a3*, we provisionally identified these as GABAergic/dopaminergic KA” interneurons (Figure 2F).

### Major neuron diversity and organization regenerate post injury

Next, we examined how these populations changed over development and regeneration. First, we found that all major neuron types (RBNs, INs, and both MNs) identified in uninjured populations were present during regeneration in our single cell dataset, indicating that neuronal diversity is largely regenerated after amputation (Figure 3A). While this confirms the presence of these neurons in the regenerating spinal cord, we next wanted to investigate if the spatial neuron organization found in uninjured tissue also regenerates. We performed Hybridization Chain Reaction (HCR) *in situ* RNA detection to visualize the expression of marker genes for major cell types (*chat-*/*prdm14+*: RBNs, *lhx1+*: INs, *chat+/prdm14-*: MNs, *chat+/prdm14+*: MNs) (Figure S3). We then manually mapped neuron borders to visualize segmented cells within the dorsal/ventral axis (Figure 3B). At the stage 41 mid-trunk, all cell types could be identified within their expected dorsal/ventral ranges (Figure 3B,C).^18^ After collecting later developmental (un3dpa, un7dpa) and regenerating (3dpa, 7dpa) samples, we found that this dorsal/ventral range did not change significantly based on condition or time, indicating that neuron spatial organization is regenerated post injury. The only exception were RBNs, which are known to degrade at later stages and could rarely be found after 3 dpa (Figure 3C).^19^ We conclude that the spatial domains of RBNs, INs, and MNs are generally re-established during regeneration, as we have previously found for neural progenitors.^23^

**Figure 3.**
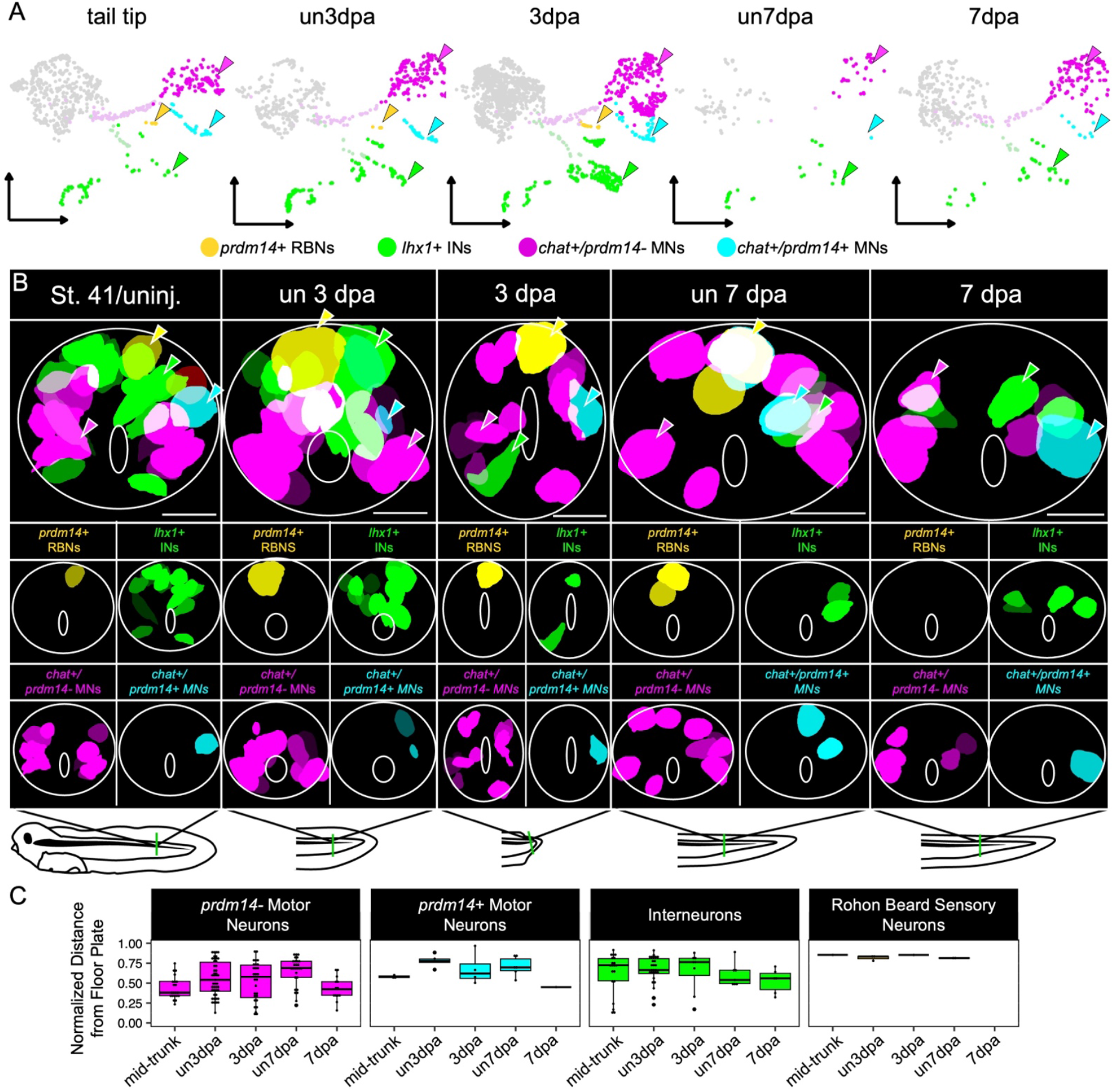
Major neuron types can be found in spatial domains within regenerating tissue. A) UMAP showing cell clusters during developmental and regenerating timepoints. B) Cell masks based on HCR fluorescence in cross-sectioned spinal cords in the stage 41 mid-trunk and in uninjured or regenerating tissue at 3 and 7 dpa. White solid line denotes spinal cord boundary (outer) and central canal (inner). Yellow is *prdm14*+ RBNs, green is *lhx1*+ INs, magenta is *chat*+ MNs, and cyan is *chat*+/*prdm14*+ MNs. Example cells of each type are shown with a color matched arrowhead. Bottom diagram shows approximate location of cross-section. All scale bars are 100 µm. C) Bar chart showing normalized distance from the floor plate for each cell type. All samples are not significantly different. St. 41/uninj n = 2 sections; un 3 dpa n = 5 sections; 3 dpa n = 9 sections; un 7 dpa n = 4 sections; 7 dpa n = 3 sections. Positive sections were selected from n = 4 tadpoles per timepoint.

### Neuron repopulation strength post-injury depends on neuronal birthdate

After confirming the regeneration of major neuron types and their spatial organization, we next wanted to see if neurons made at different times in development regenerate to the same extent in our scRNA-seq dataset. First, we defined neurons by their developmental window of neurogenesis. Spinal cord neurons have been traditionally categorized into two groups based on their time of birth during early primary or later secondary neurogenesis. To factor in additional nuance, we used the terms “early” to indicate neurons established by late neurula, “mid” for secondary neurons arising after neurulation but no longer being generated by stage 41, and “late” for neurons still undergoing neurogenesis by stage 41 (Figure 4A). We assigned a preliminary primary or secondary origin to neuron categories based on what is known from *X. laevis* and zebrafish (Figure 4B, Table S1),^18,42–45^ then confirmed this identity based on the developmental population dynamic data (Figure 4C). We found that during our developmental timepoints, primary early neurons decreased in proportion (RBNs, V2b INs), while secondary or late neurons increased in proportion beginning either at day 0 (MNs, V0v INs, KA’s) or day 1 (V2a INs, V1 INs) (Figure 4C). We identified secondary mid neurons (KA”s, dI6 INs) based on previous evidence indicating a secondary origin^14,15^ but a decrease in proportion in our dataset. Interestingly, in comparison to KA”, KA’ neurons increase during our sampled developmental stages, illustrating a novel divergence between the two cell types (Figure 4C).

**Figure 4.**
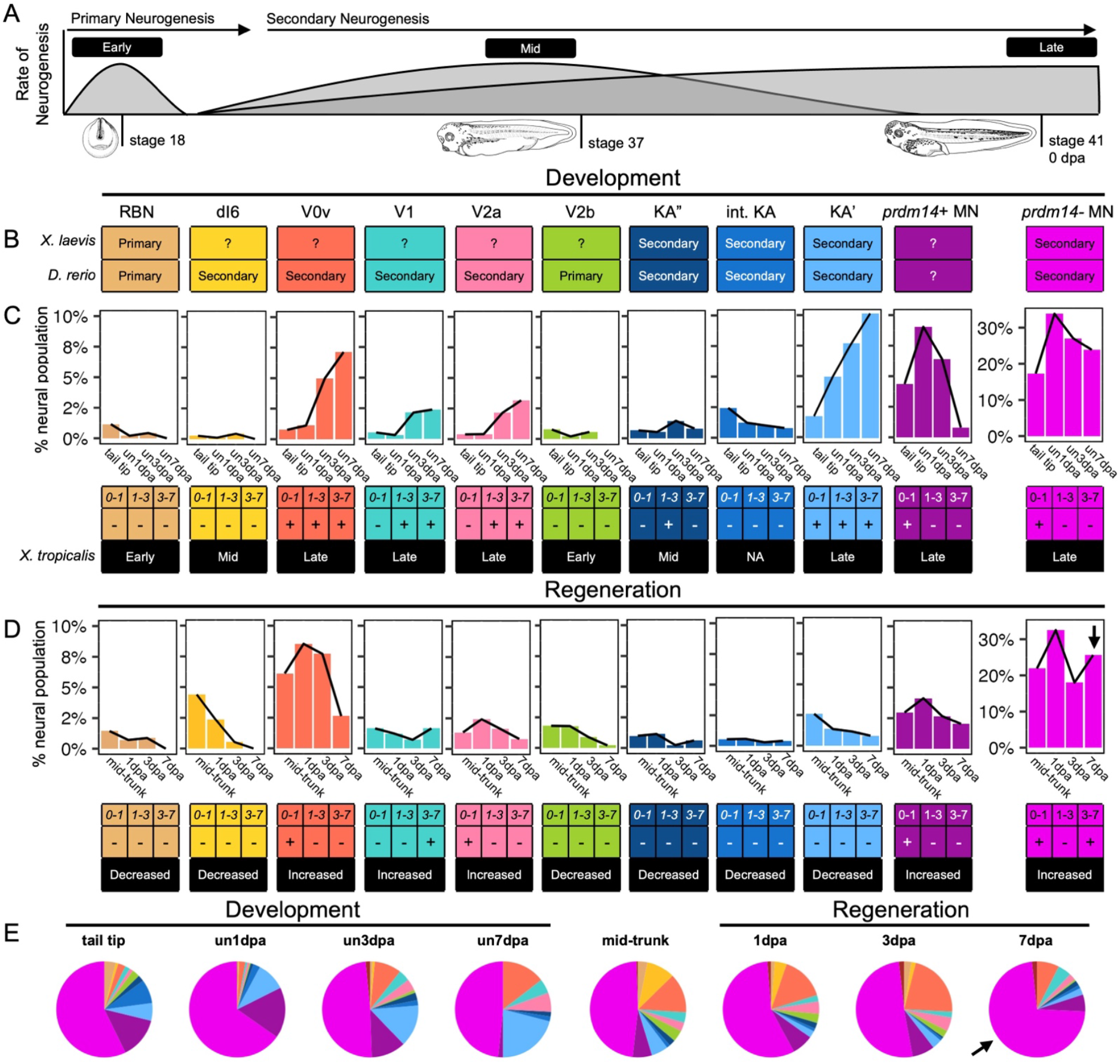
Neuron subtype repopulation after injury depends on developmental identity. A) Schematic showing waves of primary and secondary neurogenesis (y axis) over developmental time (developmental stages over x axis), additionally annotated as early neurogenesis (occurring during neurulation), mid neurogenesis (occurring between neurulation and stage 41) and late neurogenesis (occurring during stage 41). B) Literature analysis from *Xenopus laevis* or *Danio rerio* (Table S1). C) Proportional bar charts showing changes in the % neural population over developmental time, as labeled as decreased (-) or increased (+). D) Proportional bar charts per neuron type showing changes in the % neural population over regenerative time, as labeled as decreased (-) or increased (+). Arrow denotes expanded *prdm14-* MN cluster at 7dpa. E) Pie charts showing the proportion of neurons at each time point. Arrow denotes expanded *prdm14-* MN cluster at 7dpa.

If spinal cord regeneration fully recapitulates development, we would expect all neuron types to be newly generated in parallel from NPCs. Instead, neuron populations changed dynamically during regeneration based on their window of neurogenesis and identity. Most early and mid neurons, which have exited their neurogenic window by the time of amputation, proportionally decreased after injury. Most late neurons, which are still being generated at stage 41, proportionally increased during at least one regenerating timepoint (Figure 4D). However, several of these late neuron types diverged from their developmental trajectory over regenerative time. The most striking were the late *prdm14*+ MNs, V0v INs, and V2A INs, which expanded immediately after injury but then decreased markedly at later timepoints, and the late KA” neurons, which declined steadily over regenerative time (Figure 4D). Only two neuron types (*prdm14*- MNs and V1 INs) increased in proportion from 3-7 dpa. These *prdm14-* MNs undergo a unique and dynamic pattern: first expanding by 1 dpa, then declining at 3 dpa, and expanding again by 7 dpa (Figure 4D). To assess if these proportions were differentially affected by changes in the progenitor cluster we looked at internal neuron proportions (Figure 4E), which showed the universal declines seen at 3 dpa is likely due to progenitor growth, but there remains a strong expansion of *prdm14*- MNs at 1 and 7 dpa in both analyses (Figure 4D, E arrows). Overall this indicates that neurons do not universally regenerate, but instead dynamically repopulate the regenerating spinal cord depending on their neurogenic window and identity.

### Proliferative neurogenesis explains the increase in neurons from 3-7 dpa but not 0-1 dpa

The dynamic and cell-type specific changes in neuronal subtypes we observed suggested waves of neurogenesis might be occurring between 0-1 dpa and 3-7 dpa, and so we next set out to investigate if these could be attributed to proliferative neurogenesis by assessing cell cycle and Gene Ontology (GO) term enrichment.^46,47^ To assess the balance of progenitors and post-mitotic cells, we calculated the percent population of cells in S, G2M, or G1 phase using Seurat-based cell cycle prediction analysis. During development, the proportion of neural cells in G2M declines steadily from 0-7 days. This is accompanied by a decrease in NPCs and in GO:0051301 cell division score (Figure 5A-C, left panels, Figure S4). By contrast, in regeneration there is a marked increase in G2M at 3 dpa accompanied by an expansion of NPCs and an increase in the cell division score at the same timepoint (Figure 5A-C, right panels, Figure S4). At 1 dpa, the proportion of NPCs actually decreases, as does the proportion of cells in G2M, in agreement with our previous findings (Figure 5A, B).^30^ To assess if the neuron repopulation that we see from 0-1 dpa and 3-7 dpa is the result of neurogenesis, we looked for top GO terms during regeneration. Unexpectedly, we did not see an enrichment for GO:0030182 neuron differentiation terms at 1 dpa, only at 7 dpa (Figure 5D, Figure S4C, D).

**Figure 5.**
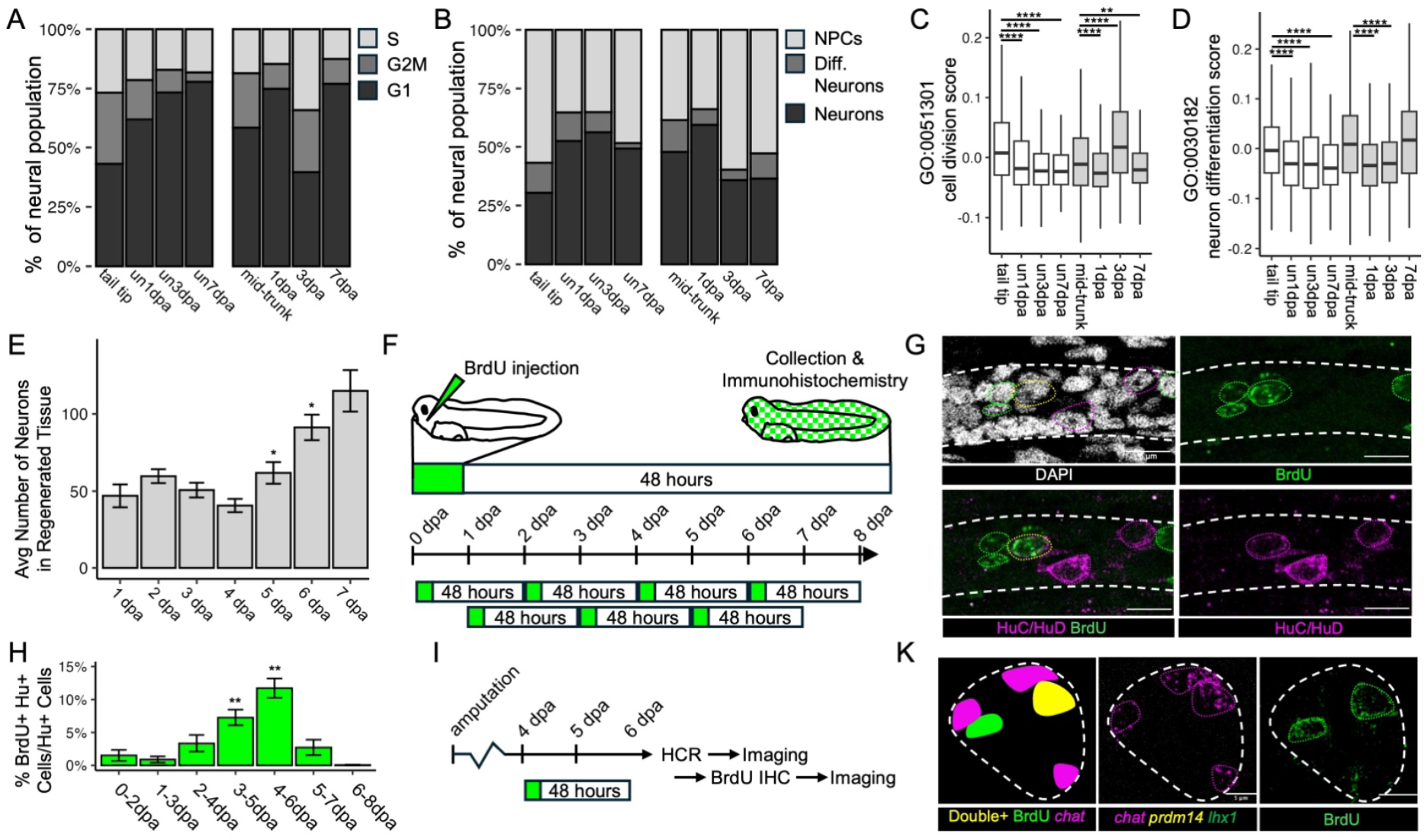
Regeneration proceeds through a wave of proliferation and then proliferative neurogenesis. A) Bar chart of G1, G2M and S phase cells as a proportion of entire neural population. B) Bar chart of NPCs, Differentiating (Diff.) Neurons, and Neurons as a proportion of entire neural population. C) GO:0051301 cell division score over developmental and regenerative time. ****p-value < 0.0001, **p-value < 0.01 compared to tail tip for developmental timepoints and mid-trunk for regenerating timepoints. D) GO:0030182 neuron differentiation score over development and regenerative time. ****p-value < 0.0001 compared to tail tip for developmental timepoints and mid-trunk for regenerating timepoints. E) Bar chart of the average number of HuC/HuD+ neurons in regenerated tissue over regeneration. * p-value < 0.05 compared to 1dpa by 1dpa n = 4; 2dpa n = 14; 3dpa n = 11; 4dpa n = 10; 5dpa n = 12; 6dpa n = 12; 7dpa = 14. F) Experimental diagram of BrdU injection and BrdU/HuC/HuD Immunohistochemistry. G) Whole mount image showing double HuC/HuD and BrdU+ cell at 6 dpa after injection at 4 dpa. White dashed lines indicate spinal cord border; magenta dotted lines indicate HuC/HuD+ neurons; green dotted lines indicate BrdU+ cells; yellow dotted line indicates double positive cell. Scale bar = 5 µm. H) Plot showing the percent of BrdU/HuC/HuD+ cells within HuC/HuD+ cells 48 hours post injection. ** p-value < 0.01 compared to 0-2 dpa timepoint by . 0d-2d n = 6; 1d-3d n = 7; 2d-4d n = 6; 3d-5d n = 5; 4d-6d n = 7; 5d-7d n = 6; 6d-8d n = 7. I) Experimental diagram showing BrdU injection, sample collection and iterative co-staining with HCR and BrdU. K) Cross-section image showing *chat*+/*prdm14-* BrdU+ MN at 6 dpa after injection at 4 dpa. White dashed lines indicate spinal cord border; green dotted lines indicate BrdU+ Neurons; magenta dotted lines indicate chat+/prdm14-cells; yellow dotted line indicates double positive cell. Scale bar = 5 µm. n = 3 *prdm14*+BrdU+ cells from 10 sections generated from 4 tadpoles. All statistical tests done via Wilcoxon test.

This analysis suggested that only the expansion of V1 INs and *prdm14-* MNs we see from 3-7 dpa could be explained by new differentiation of neurons from mitotically active progenitors, or proliferative neurogenesis. To confirm these results *in vivo*, we tracked neuron repopulation by staining for the pan-neuronal marker HuC/HuD during regeneration. We found that starting at 1 dpa, the average number of neurons per regenerate does not increase until 5 dpa, where it increases until our endpoint at 7 dpa (Figure 5E). To see if this increase in neuron population at 5 dpa is due to proliferative neurogenesis, we injected tadpoles with BrdU according to established protocols.^48^ A 2-day chase was necessary after injection to detect substantial neurogenesis (Figure 5F). We found that there is little to no proliferative neurogenesis in the regenerated tissue prior to 3 dpa, as less than 5% of neurons were BrdU+. Instead, there is a peak in neurogenesis at 4-6 dpa, with ∼12% double positive cells, before declining to baseline levels (Figure 5G, H). We conclude that proliferative neurogenesis begins between 3 dpa and 4 dpa, which is reflected in the increase of NPCs, cells in G2M, and cell division scores at 3dpa (Figure 5A, B, C). This likely fuels the formation of new neurons we observe 5-7 dpa (Figure 5E), as well as the increases in Differentiating Neurons and GO neuron differentiation score we observe at 7 dpa (Figure 5B, D).

To test which neuron types were being generated by proliferative neurogenesis, we did iterative staining of HCR and BrdU (Figure 5I). The only double positive cells we found at 7 dpa were BrdU+ *chat*+/*prdm14*- cells, indicating that the most common BrdU+ HuC/HuD+ neurons generated 5-7 dpa are either V2a INs or *prdm14*- MNs (Figure 5K), which echoes the strong increase in *prdm14*- MNs at 7 dpa compared to most other neuron types (Figure 4). However, proliferative neurogenesis cannot explain the increase in neurons observed at 1dpa (Figure 5B) or the increase in V0v, V2a, *prdm14+* and *prdm14-* MNs at that timepoint (Figure 4D) (see Discussion).

### Emergence of transient population of regeneration-specific neurons at 1dpa

In zebrafish, a transient group of injury induced neurons (iNeurons) arises soon after spinal cord injury and plays a central role in cell-cell signaling and spontaneous neuronal plasticity and thus, functional recovery. iNeuron markers were shown to colocalize with various neuron markers such as *hb9, isl1, pax2, gad1b, and vgluta*, indicating they arise from various neuron types after injury.^27^ Given the lack of both proliferation and neurogenesis at 1 dpa as well as previous evidence of a transient class of *lepr*+ regeneration-specific neurons in *Xenopus*,^30,36^ we hypothesized that pro-regenerative mechanisms are enacted by a similar population of injury-responsive neurons at this timepoint.

To test this hypothesis, we identified subclusters of neurons that appeared transiently after injury (Figure 6A). These transient subclusters arose in V2a INs, *prdm14*+ MNs, *prdm14*- MNs, V0v INs, and RBNs primarily at 1 and 3 dpa (Figure 6B). Despite this temporal commonality, each subcluster showed unique patterns of gene expression. In comparison to the RBN and V0v regeneration-specific subclusters, V2a IN, *prdm14*- MN, and *prdm14*+ MN subclusters all strongly expressed *atf3*, a gene previously implicated in neuron regeneration and a marker of zebrafish iNeurons (Figure 6C).^27,49^ This supports previous work showing that transient regeneration specific neurons do not represent one neuron type, but rather arise within several types.^27^ The *atf3*+ MN subclusters also express leptin receptor (*lepr*), which has been previously identified in a regeneration-specific MN cluster during neuron regeneration in *Xenopus*,^30,36,50^ indicating that zebrafish iNeurons and *Xenopus lepr*+ transient MNs may represent a conserved regeneration-specific mechanism (Figure 6C).

**Figure 6.**
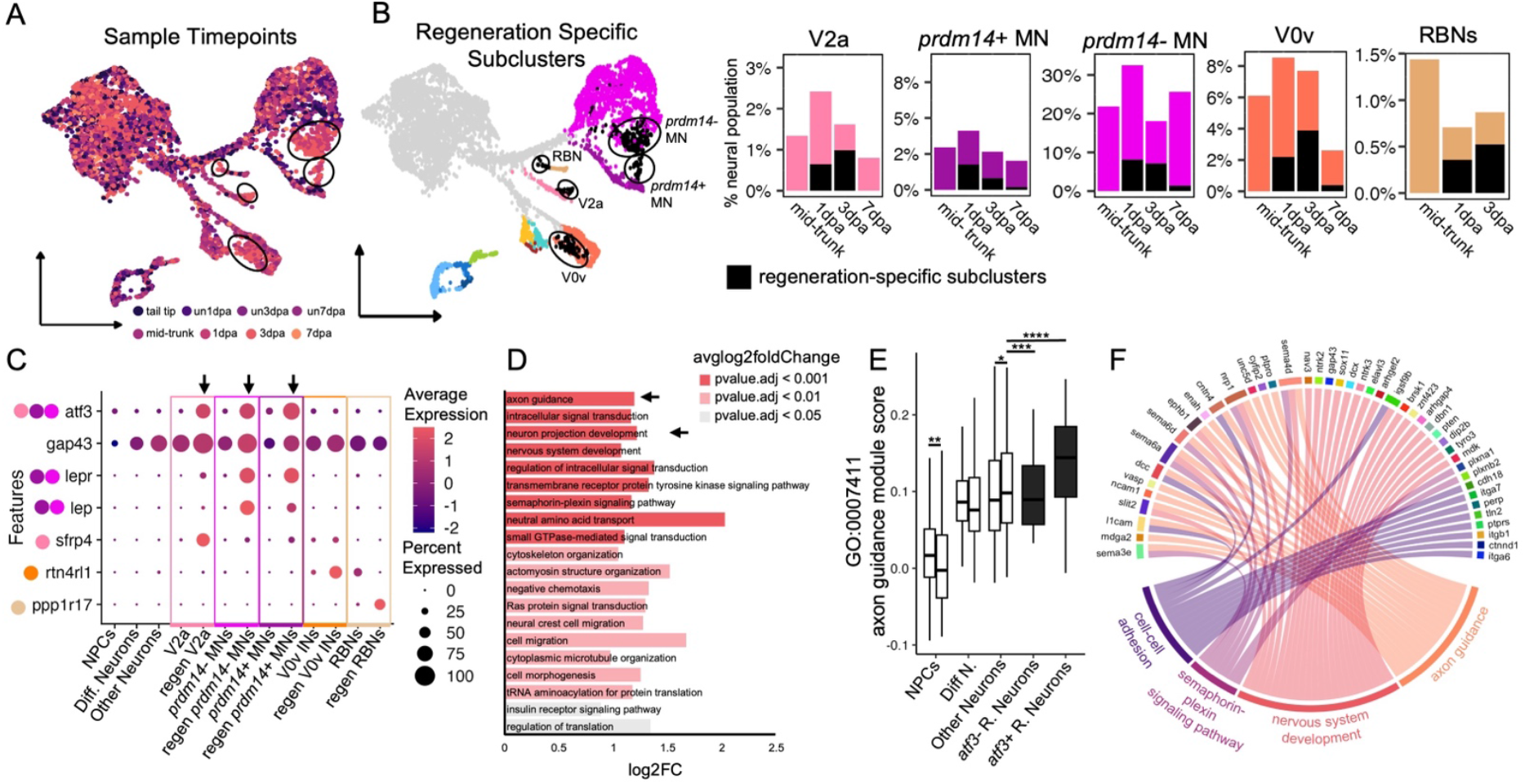
Regeneration-specific neuron subclusters arise in specific neuron types and show distinct patterns of gene expression and cell processes. A) Combined UMAP showing sample distribution; black circles show regeneration-specific subclusters. B) Regeneration specific subclusters (black) appear in V2a INs, *prdm14*+ MNs, *prdm14-* MNs, V0v INs, and RBNs by 1 dpa, as shown by combined UMAP (left) and proportional bar charts (right). C) Gene expression of NPCs, Differentiating (Diff) Neurons, Other Neuron types, and regeneration-specific subclusters. Some subclusters express iNeuron markers (*atf3, gap43*), others express *Xenopus lep*+ MN markers (*lepr, lep*), and others have unique expression patterns (*sfrp4, rtn4rl1, ppp1r17*). Arrows highlight *atf3*+ Regenerating Neurons. Y-axis dots represent clusters gene is expressed in. D) Gene ontology of *atf3*+ Regenerating Neurons compared to other Neuron types at 1dpa. Enriched terms are sorted by strength of adjusted p-value. E) Plot of GO: 0007411 axon guidance score at 1 dpa in different cell clusters in uninjured (left bar) or regenerating (right bar) timepoints. Black denotes regeneration-specific subclusters. *pvalue < 0.05, **pvalue < 0.01, ***pvalue < 0.001, ****pvalue < 0.0001 F) Plot of GO:0007411 axon guidance score and associated modules at 1 dpa (bottom), compared to upregulated genes representing scores. All statistical tests done via Wilcoxon test.

Previously, zebrafish iNeurons were found to have a role in cell signaling and neural plasticity through cell morphogenetic mechanisms like axon regrowth and migration.^27^ To assess if *atf3*+ subclusters play a similar central signaling role in *Xenopus*, we used CellChat to find top signaling pathways.^51^ Overall, while many top pathways were similar to iNeurons (ex/ L1CAM signaling), the differences in total signaling strength between regenerating neurons and non-regeneration specific clusters was insignificant (Figure S5). To then interrogate if *atf3*+ subclusters drive spontaneous neuronal plasticity, we investigated what GO terms were enriched in *atf3*+ regenerating neurons compared to non-regeneration specific neurons at 1 dpa. Similar to zebrafish iNeurons, the top enriched GO term was axon guidance, with other top terms including intracellular signaling transduction (Figure S6A, B), neuron projection development (also increased in zebrafish iNeurons), and nervous system development (Figure 6D, E, S6). To see what genes were driving this enrichment, we plotted genes associated with nervous system development against axon guidance and other top overlapping associated GO terms. Many nervous system development genes were associated with axon guidance and specifically the semaphorin-plexin signaling pathway and/or cell-cell adhesion (Figure 6F), indicating that like iNeurons, *atf3*+ cells prioritize morphogenetic plasticity during early regeneration (Figure S6D-E).

### Movement of anterior neurons contributes to repopulation of the regenerating tail

Though all neuron types exist in regenerated tissue, proliferative neurogenesis doesn’t occur until 4 dpa (Figure 5) and is only clearly identifiable for some neuron types. This raises the question of how other neurons appear in new tissue without new neuron differentiation. A possible mechanism is movement of post-mitotic, differentiated neurons from uninjured tissue into new tissue, as has been reported in urodeles.^23^ We tested for neuron movement via two methods, focusing on KA” INs, because they are present in two ventral columns with well-spaced cell soma and can be visualized with an antibody against TH. First, we quantified the number of dopaminergic/GABAergic KA” INs in pre-amputation, un7dpa, and 7 dpa tails to test if the total number of neurons are retained or if more are generated during regeneration. We found that prior to amputation, an average of 61 TH+ KA” INs populate the spinal cord from vent to amputation plane. After 7 days, un7dpa samples have developed on average 125 TH+ KA” INs, likely due to maturation of existing cells. By contrast, regenerating 7dpa samples do not recover lost density and instead retain on average only 70 KA” neurons across the whole spinal cord, with ∼12 within the regenerating portion (Figure 7A). To confirm that these cells were not a result of proliferative neurogenesis, we injected tadpoles with BrdU at 4 dpa before collection at 6 dpa and co-staining with TH. All TH+ KA” INs identified were BrdU- (Figure 7B). To see what cell processes were occurring in KA” INs at this time, we looked at GO terms and found enrichment for cell migration in several cells, including KA” INs (Figure S7). To test if we could inhibit neuron movement by blocking cell migration, we inhibited migration factor *dclk* and counted TH+ KA” INs, finding a decrease in the number of KA” INs in the regenerating spinal cord (Figure S7).

**Figure 7.**
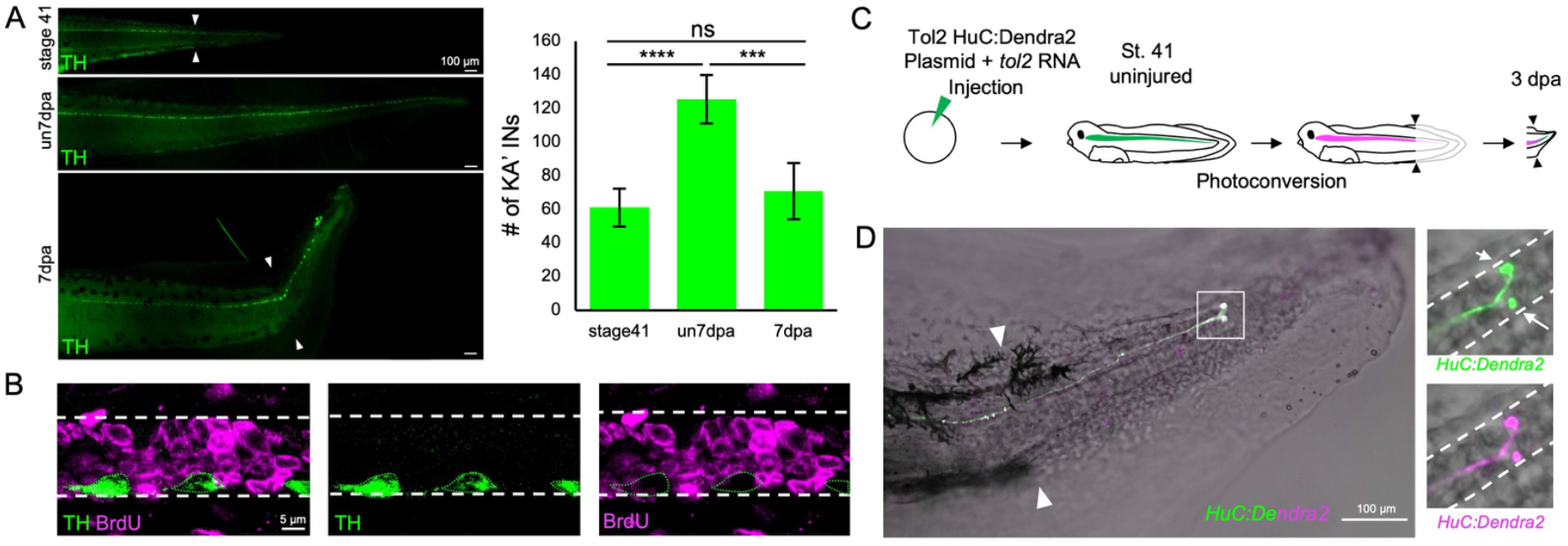
Post-mitotic neurons are displaced from uninjured tissue to regenerating tissue. A) TH+ KA” INs in the uninjured stage 41 (0 dpa) or un 7 dpa tadpole tail compared to the 7 dpa regenerate; paired triangles represent amputation plane (left). Bar chart showing the average number of TH+ KA” INs (n = 4 per timepoint) (right). B) Image of 6 dpa tadpole co-stained for TH and BrdU. Green dotted lines represent TH+ KA” INs, white dashed lines show spinal cord border. C) Diagram of experimental design: Tol2 HuC:Dendra2 plasmid and *tol2* RNA was injected into the single-celled embryo. At stage 41, tadpoles were photoconverted and amputated. At 3 dpa, tadpoles were imaged. B) Representative image of double-positive neuron in the regenerated tail. Paired triangles represent amputation plane. White box represents insets of the 488 (top, green) and 594 (bottom, magenta) channels. White dashed lines represent spinal cord border.

To directly confirm that mature neurons were displaced from uninjured tissue, we injected 1-cell stage embryos with *tol2* mRNA and a donor plasmid containing sequence for the photoconvertible protein Dendra2 under the control of the mouse Hu promoter. Dendra2 is subsequently expressed mosaically in a small fraction of Hu+ neurons. At stage 41, the entire tail was photoconverted, amputated, and tadpole allowed to regenerate (Figure 7C). Any neurons present before the injury would thus appear double positive, while neurons generated subsequently would appear green. At 3 dpa, we found double positive Dendra2 cells in the regenerating spinal cord, confirming that neurons move from uninjured tissue into the regenerating spinal cord (Figure 7D).

## Discussion

Our study creates a molecular atlas of neurons in the *Xenopus* tadpole, defining marker genes for both known and previously unconfirmed neuronal types. We establish that the overall spatial organization of regeneration largely recapitulates developmental organization, but that the cell type composition of the regenerating spinal cord differs in many respects from its uninjured, developing counterpart. Proliferative neurogenesis occurs both less and later than expected, giving rise principally to *prdm14*- MNs. Some but not all neuron types give rise to a transient population of neurons that prioritize morphogenetic plasticity as early as 1 dpa. Other neuron types, including KA” INs, repopulate the spinal cord without proliferation, implicating post-mitotic movement (Figure 8). These findings have important implications for understanding the mechanisms that may enable plasticity in regenerative conditions, or limit repair in non-regenerative conditions.

**Figure 8.**
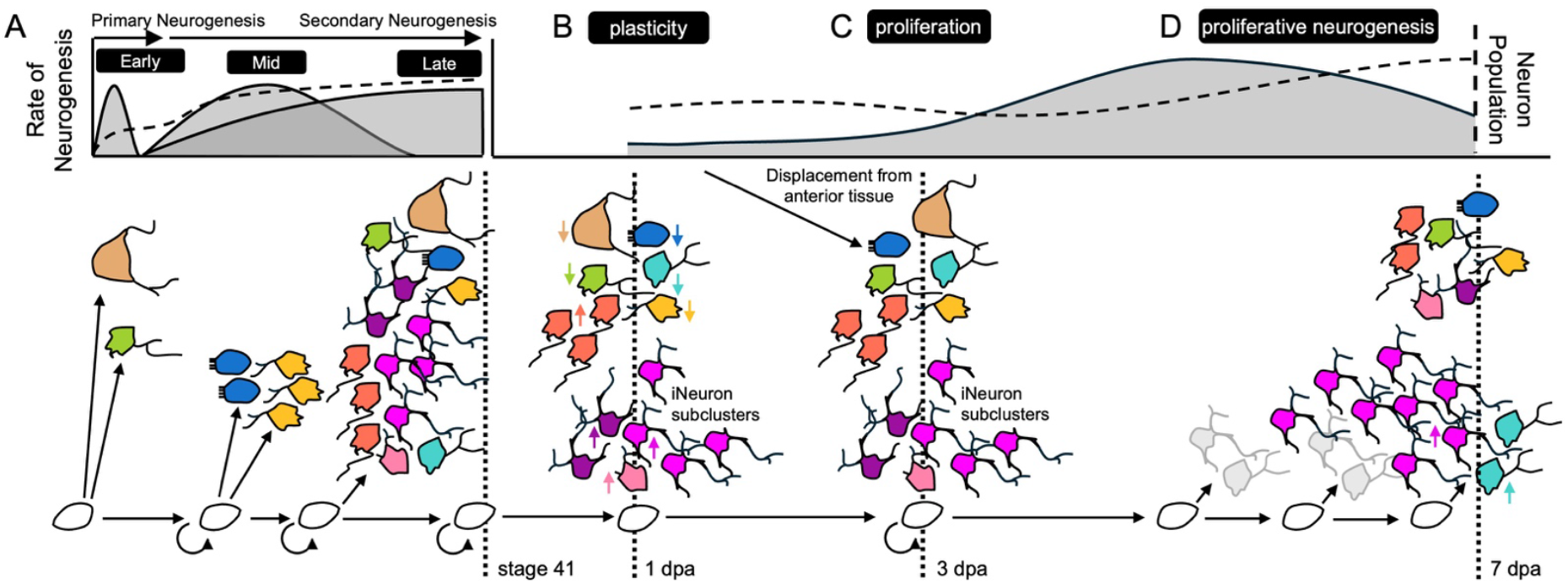
Neuron regeneration proceeds dependent on neuron identity. A) During development, primary and secondary neurogenesis gives rise to early, mid, and late neuron types, respectively, resulting in a complex population of neurons by stage 41. B) At 1 dpa, most late neurons increase in proportion in comparison to early and late neurons. Putative iNeuron clusters arise from *prdm14*+ MNs, *prdm14-* MNs, and V2a INs and drive neural plasticity. C) At 3 dpa, NPCs prioritize proliferation, although putative iNeuron clusters remain present. Few new neurons are generated, but neurons from uninjured tissue move into regenerating tissue. D) New neurons—primarily *prdm14-* MNs—are generated by proliferative neurogenesis.

### A molecular atlas of neuronal subtypes in the free-swimming tadpole

One of our principal goals was to better index the neuron types that exist in the early *Xenopus* free-swimming tadpole. By stage 41, tadpole motor circuits for locomotion and escape must be fully functional, but molecular identifiers for all types of neuron populations were sparse.^17^ We were able to confirm the presence of and identify markers for each major neuron type and seven conserved cardinal classes of interneurons that have previously been identified in zebrafish and mouse.^52^ In addition, we identified several novel subtypes; we were especially interested to find that there are two molecularly distinct populations of Kolmer-Agduhr neurons that could be defined by gene markers and GABAergic/dopaminergic vs GABAergic gene expression. Based on these conserved markers, these populations align to the KA’ and KA” populations found in zebrafish and mice.^41^ However, no prior reports of dopaminergic activity in KA interneurons have been reported in *Xenopus* or zebrafish, although serotonin expression has been reported in certain types of KA neurons in fish.^53^ In addition, we found two novel subpopulations of MNs (*prdm14*- and *prdm14+)*, although the functional distinction between these populations remains unclear. Although patterns of secondary neurogenesis in the tadpole were not previously established for all the cell types we found, our observations agree with what is known in *Xenopus laevis* and show parallels to zebrafish.^18,42–45^ Cell types thought to be established through primary neurogenesis, including RBNs and V2b interneurons, remain largely static in number during the 7 days of development we captured (from stage 41-47). By contrast, neuron types that arise through secondary neurogenesis, including V0v, V1, and V2a INs, expand steadily during this time.

### Similarities and differences between the uninjured developing spinal cord and regeneration

A common hypothesis is that regeneration in developing organisms will completely recapitulate developmental mechanisms to replace lost tissue.^54^ Our study supports some aspects of this model. We find the spatial organization of the spinal cord is generally restored after injury. We find that all neuron types are present in the regenerating spinal cord, however, neurons still undergoing secondary neurogenesis at stage 41 regenerate much more abundantly than those known to be established through primary neurogenesis. This may reflect the ability of the larval spinal cord to exploit the same growth and patterning mechanisms already in place before injury to readily replace neurons that are still being generated developmentally at that stage. However, the composition and dynamics of neuron types in the regenerating spinal cord differ in several crucial aspects from their uninjured, developing, counterparts. Among secondary neurogenic cell types, there are notable differences in their dynamics during regeneration versus development. Several cell types that expand during development (V0, V1, V2a, and KA” INs) instead decline in late regeneration, compensated by non-developmental increases in *prdm14*- MNs. Overall, this indicates that regenerative repopulation depends closely on developmental identity: early primary and mid secondary neurons are no longer able to dynamically repopulate in comparison to late secondary neurons, which are already undergoing population increases during development at these stages. This illustrates a divergence from the model where development is a complete recapitulation of developmental mechanisms to restore lost tissue complexity.

### Neurogenic strategies vary by cell type and temporal window

Another major motivation for our study was to understand how different classes of neurons are restored over the course of regeneration. Here we found that there are several neurogenic strategies employed that vary by time and cell type, and that some classes of neurons only form after injury. Previously, we had described that NPCs undergo an early cell cycle exit after injury, though we weren’t able to answer the question of if and when proliferative neurogenesis occured^30^. Here, the increased duration of our study and the inclusion of stage-matched developmental timepoints allowed us to generate a much clearer picture of how neurogenesis proceeds after injury. We find that proliferative neurogenesis certainly does contribute to spinal cord regeneration, but not until intermediate regeneration, with BrdU incorporation into neurons first identifiable at 3 dpa.

Proliferative neurogenesis can be tied to the late expansion we see in *prdm14*- MNs, with BrdU detectable in *chat+ prdm14-* neurons. We confirmed previous findings showing the emergence of regeneration-specific subclusters that each show unique patterns of gene expression: subclusters arise in both MN groups and V2a INs that express zebrafish iNeuron markers, and the MN subclusters additionally express *lep/lepr*.^27^ Strikingly, expression of these markers was found in several but not all neuronal classes, suggesting that some neuron types are primed to adapt post-injury. This seems to only be selectively correlated with birth time: no primary neuron classes expressed *atf3* post-injury, but several secondary classes also did not express *atf3*. It is also not clearly correlated with connectivity, although the prioritization of neuronal plasticity at 1 dpa may indicate these neurons are critical to reinnervation of new tissue.

Finally, we addressed what mechanisms, if not proliferative neurogenesis, could underlie neuron repopulation. First we focused on KA” neurons as a representation of postmitotic neurons and found that the average total number of KA” INs in the spinal cord stayed the same post injury, indicating that cells found in the regenerating tissue were from anterior, uninjured tissue. This movement was confirmed by BrdU analysis and photoconversion tracing. Due to the late emergence and small percentage of BrdU+ neurons during regeneration (12% at the peak of proliferative neurogenesis), it’s possible that this neuron movement represents a substantial proportion of the neuron repopulation we see at early and intermediate regenerative stages. While some evidence implicates active migration as a mechanism enabling repopulation of these neurons in the spinal cord, though we cannot rule out passive movement or direct differentiation from NPCs.

Looking forward, we are especially interested to see how the contributions of *atf3*+ neurons, neuronal migration, and proliferative neurogenesis change as the tadpole progresses through metamorphosis. The stage-specific loss of regenerative competence in the spinal cord is a unique advantage of *Xenopus* for understanding how regeneration becomes limited. While age-specific losses of neural stem cells and changes in the innate immune system are known contributing factors to regenerative loss, this study highlights injury-responsive neurons and movement of anterior neurons as other events that are necessary for restoring the full neuronal diversity of the spinal cord and may be differentially lost with maturity.

## Methods

### *X. tropicalis* husbandry and use

Use of *Xenopus tropicalis* was carried out under the approval and oversight of the IACUC committee at UW, an AALAC-accredited institution. Generation of tadpoles was carried out according to published methods.^30^ In this study we used both wild-type frogs and frogs from the triple transgenic line Xtr.Tg(*pax6*:GFP;*cryga*:RFP;*actc1*:RFP) (RRID:NXR_1.0021^55,56^) reared and purchased from the National Xenopus Resource (https://www.mbl.edu/xenopus/; RRID:SCR_013731). Pax6 transgenic matings were performed by crossing a heterozygous transgenic frog to a wild-type frog. These matings yielded clutches with 50/50 wt/*pax6*:GFP+ populations. For amputation assay, NF stage 41 tadpoles were anesthetized with 0.05% MS-222 in 1/9x MR and tested for response to touch prior to amputation surgery. Once fully anesthetized, a sterilized scalpel was used to amputate the posterior third of the tail. Amputated tadpoles were removed from anesthetic media within 10 min of amputation into new 1/9x MR. Tadpoles were kept at a density of no more than 1 tadpoles per 2.5 mL. Tadpoles were fed daily sera Micron^57^ at ¼ mL per 2.5 L, starting at 3 dpa until final timepoint collection.

To measure regenerated length of tail, samples were mounted and imaged with a Leica M205FA fluorescence Stereomicroscope. Using the Leica imaging software (LasX), measurements were taken from the vent to the most posterior point of the tail. Regenerating measurements were taken based off of morphological identification of amputation plane.

### scRNA-sequencing pipeline

Tadpoles were anesthetized with 0.05% MS-222 in 1/9x MR and tested for response to touch prior to amputation surgery. Once fully anesthetized, a sterilized scalpel was used to amputate and collect intended tissue (Figure 1A). Tail tissues were collected within 30 minutes of amputation and added to a vial with 72 µl trypLE (ThermoFisher Scientific:12604013) and 3 µl 100 mg/mL collagenase P (ROCHE:11213857001) for every 100 tail tissues. Samples were moved to a 28C water bath for dissociation and triturated with a p200 every 5 minutes until dissociated fully and no chunks was visible (30-60 minutes). To quench the reaction, 150 µl of ice cold 10% FBS was added for every 100 tail tissues. Samples were then spun down for 5 minutes at 1000xg. Supernatant was disposed and cells were resuspended in 300 µl 10% FBS for FACS. For each sample, 22-294 tadpoles were used (tail tip: 294 tadpoles; un1dpa: 222 tadpoles; un3dpa: 133 tadpoles; un7dpa: 22 tadpoles; mid-trunk: 294 tadpoles; 1dpa: 208 tadpoles; 3dpa: 144 tadpoles; 7dpa: 25 tadpoles). Tail tip and mid-trunk tissues were collected from the same specimens.

Cell sorting and collecting were performed as previously described,^30^ except cells were collected into 500 µl solutions of 1x PBS with 1 mg/ml BSA (NEB:B9200S). Single-cell mRNA libraries were prepared using the GEM-X single cell 3’ v4 kit (10x Genomics), with a target capture of 10,000 cells. Quality control and quantification assays were performed using a Qubit fluorometer (Thermofisher) and a Screentape Assay (Agilent). Libraries were sequenced on an Illumina NextSeq2000 using a 100 cycle, P4 XLEAP-SBS™ kit. Each sample was sequenced to an average read depth of 226 million total reads, which resulted in an average read depth of ∼28,000 reads/cell after normalization.

### scRNA-sequencing processing

RNA sequencing reads were processed using CellRanger v9.01 Count and Aggregate by 10x Genomics. To generate the reference genome, the *Xenopus tropicalis* v10.0 reference genome and gene models were downloaded from Xenbase. The output from CellRanger was processed using DoubletFinder to detect doublets. The data was then run through the Seurat v3.1 pipeline^35^ in R Studio v4.4.3 using standard filtering and preprocessing. Cells expressing < 200 genes were removed from downstream analysis, with a mitochondrial RNA read cutoff of 25%. Two poor quality clusters indicated by abnormally low UMI count and high mitochondrial RNA percentage were manually removed. After QC analysis, cells were normalized, scaled by Sample and nFeature_RNA, then integrated with RPCA. Cell cycle was predicted as previously described.^30^

Go.db (v3.21) was used to evaluate Gene Ontology^46,47^ terms during regeneration, and circlize (v0.6.16)^58^ was used to generate chord graphs of gene-term connectivity. CellChat (v1.4.0)^51^ was used to evaluate *atf3*+ regenerating neuron cell-cell interactions. Only *Xenopus* genes with 1-to-1 ortholog mapping, as defined on Xenbase, were mapped to mouse orthologs.

### Hybridization Chain Reaction and Sectioning

For HCR, tadpoles were fixed overnight in 1x MEM with 3.7% formaldehyde at 4C. HCR was done in whole mount from established protocols,^59^ at a probe concentration of 16 nM. Post HCR protocol, samples were washed with 3 x 15 min 1X PBS, 1 x 15 min 15% sucrose in 1X PBS, then overnight at 4°C. To identify regenerating tissue post-sectioning, fins were trimmed dorsal and ventrally only posterior to the amputation plane. Samples were then transferred into OCT (Thermofisher 23-750-571) and kept at -80°C until sectioning with a Leica CM3050 S Cryostat. Post sectioning, slides were baked at 37C for 20 minutes before imaging using a Leica SP8 Confocal with a 63X objective. They were processed using FIJI image analysis software. For cell masks, Cellpose^60^ was manually used to create 3D cell masks based on HCR and DAPI staining that were then imported into FIJI for downstream processing. To measure Dorsal/Ventral location, the center point of cells was measured from the bottom of the spinal cord floor plate then normalized to spinal cord height, as measured from the bottom of the floor plate to the top of the roof plate.

### BrdU Labeling

In order to assess the level of proliferative neurogenesis during regeneration, we used the DNA-synthesis marker BrdU. Tadpoles at the indicated stage were injected with 16 nL of BrdU (10 mM, Fisher, B23151) in the gills, following the protocol established *Xenopus* for tracking neurogenesis during development.^48^ After injection, tadpoles were allowed to regenerate for 2 days to allow for detectable differentiation into neurons via HuC/D staining. We found labeling in the spinal cord within 1 hour after injection and that labeling persists in progenitors for ∼2 days post-injection (data not shown). Staining proceeded as previously described,^23^ (1° Abs: 1:200 Rb anti-BrdU polyclonal, Fisher, PA5-32256; 1:200 Ms HuC/D monoclonal, Fisher, A21271) with the addition of a 10 minute 2N HCl acid wash following the initial permeabilization in PBS-Triton. Because the BrdU 10 minute 2N HCl wash destroys HCR signal, we iteratively stained for HCR and BrdU. Samples were injected with BrdU as described then run through the HCR protocol, sectioned and imaged as described above. Slides then underwent staining for BrdU and re-imaging using the described protocol, with solution volumes adjusted for slide Immunohistochemistry.

### TH Immunohistochemistry

Tadpoles were stained and imaged as previously described,^23^ (1° Abs: 1:200 M anti-Tyrosine Hydroxylase monoclonal, Immunostar, 22941). Co-staining with BrdU was conducted as described above.

### Dclk1 inhibitor treatments

Tadpoles were treated by immersion in 5µM Dclk1-in1 (Tocris cat #7285; 50mM stock solution was made up in DMSO, diluent for immersion was 1/9MR) either immediately following amputation or at the corresponding uninjured timepoint. 0.01% DMSO was used as a vehicle control. Inhibitor solution was refreshed on days 1, 3, and 5.

### Dendra2 Photoconversion

In order to track migration and differentiate between newly generated neuronal cells, we used a pMT-HuC: Dendra2 plasmid on post-mitotic neurons. The tadpoles were injected at the one-cell and two-cell embryo stages with 2 nanoliters of Dendra2 plasmid. After injection, *Xenopus Tropicalis* tadpoles were grown to stage 41 where we took pictures using the 20X objective on a compound microscope. Then we conducted photoconversions for differencing between old and new neurons by directing UV light on a posterior section of the tail. Then the distal 1/3 of the tail was amputated using a scalpel. At 3 days post-amputation (dpa), the tails were once again imaged in the 488 nm and 594 nm channels.

### Statistics

All statistics were performed in using R Stats package (v3.6.2).

## Acknowledgements

We thank members of the Wills lab for critical comments during the preparation of this manuscript and support in frog husbandry. We thank the ISCRM Genomics Core and Director Mary Regier, for equipment use and their support during the scRNA-Seq pipeline. We thank the UW Cell Analysis Facility for the equipment use and their support for cell sorting. We thank the Ruohola-Baker lab for the use of their equipment for tissue processing during cell dissociation. We thank the Kong lab for access and training on their cryostats. We thank Xenbase for curation of genomic and literature information and the National Xenopus Resource for frogs. AAS was supported by the Cellular and Molecular Biology Training Grant NRSA 5T32GM136534-02 from NIGMS. SBP was supported by a Mary Gates Fellowship for Undergraduate Research. This work was supported by NINDS R01NS099124 to AEW.

## Author Contributions

AAS and AEW conceived of the idea. AAS performed the 10X pipeline and conducted Seurat analysis with support from SBP. AAS developed and performed HCR experiments. SBP developed and performed BrdU experiments. IH and TAF performed Dendra2 photoconversion experiments. MEM performed control TH staining experiment. AAS and AEW wrote the manuscript with support from SBP.

