## Supplemental Figures for "Spinal cord regeneration deploys cell-type specific developmental and non-developmental strategies to restore neuron diversity"

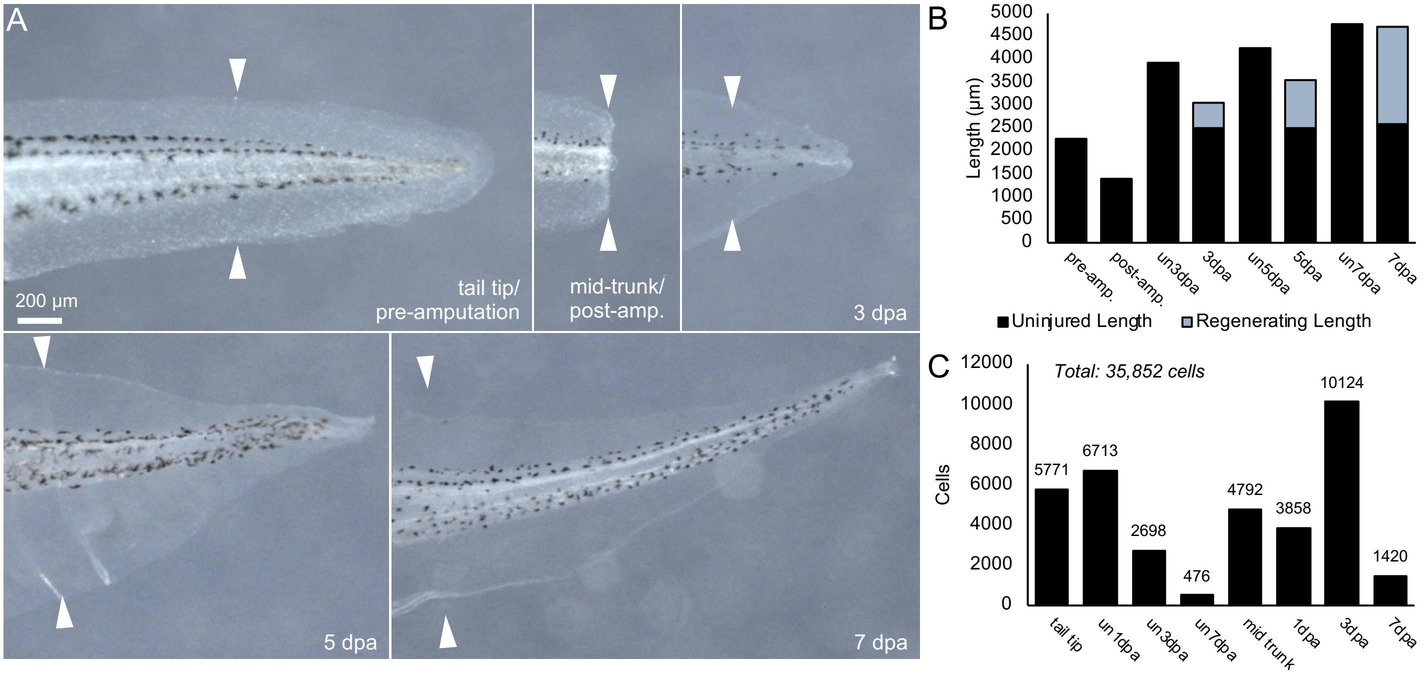


**Figure S1. Experimental 7 day time course of regenerating stage 41 tadpoles.** A) Brightfield images of tadpoles at tail tip/pre-amputation (pre-amp.) stage 41, mid-trunk/post-amputation (post-amp.), 3 dpa, 5 dpa, and 7 dpa. B) Bar chart of average vent-to-tip tail length in microns of uninjured (un-) or regenerating tails. Black denotes uninjured tissue, grey denotes regenerated tissue. C) Bar chart of cells collected by scRNA-seq, post quality control analysis; numbers above bars show cell counts per timepoint, total dataset is 35,852 cells.

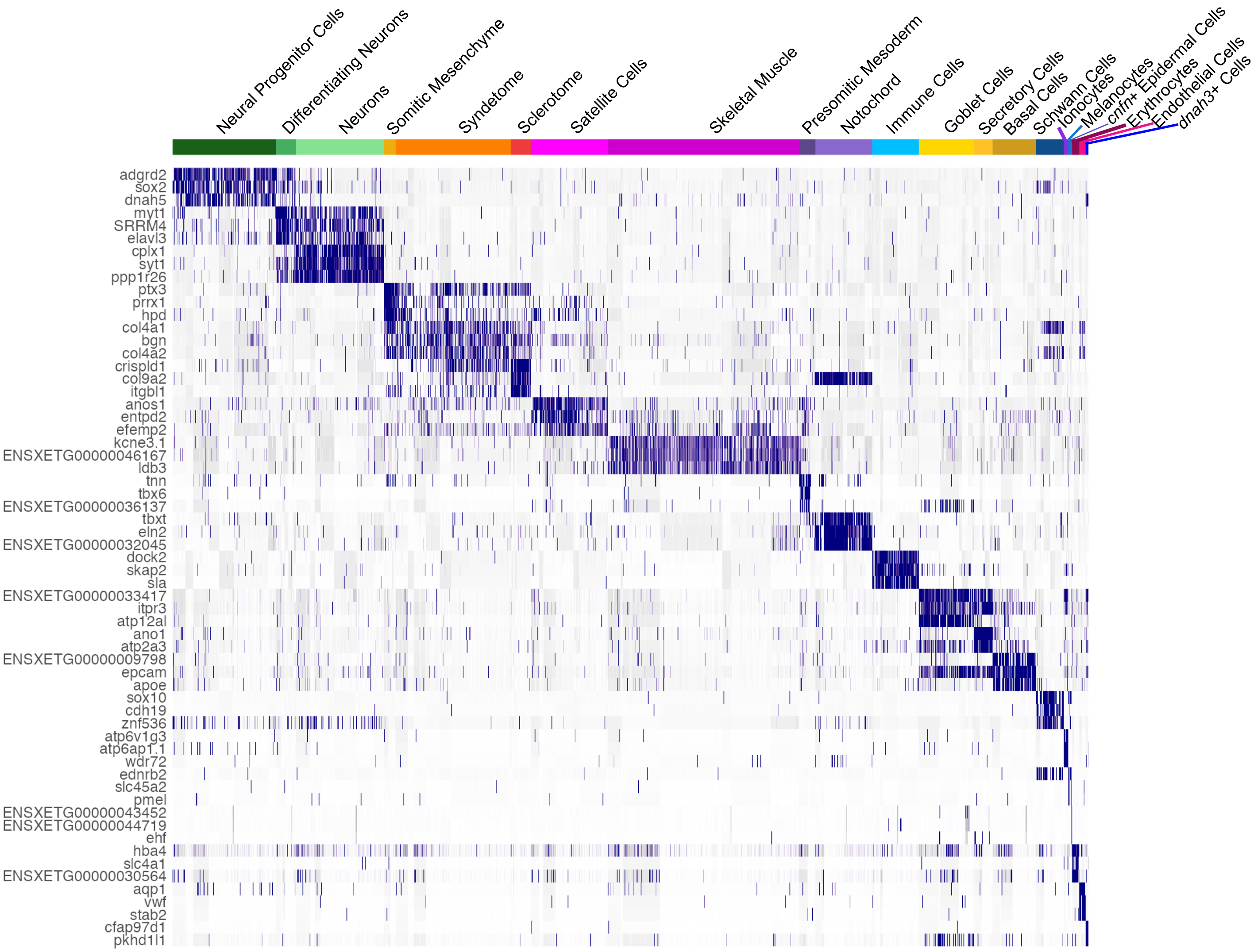


**Figure S2. Gene expression patterns identify tail cell types.** Heatmap of top three differentially expressed genes for each identified cell type. Purple indicates high expression, white indicates low expression.

| **Cardinal Classification** | **Neurotransmitter** | **Zebrafish** | **likely *Xenopus* counterpart** | **Birth Date** | **Marker Genes** | **Potential Subcluster Notes** |
|---|---|---|---|---|---|---|
| RBN | Glutamatergic | Rohon Beard Neurons<br>- Tuttle et al., 2024, | Rohon Beard Neurons | Primary | *prdm14, isl2, isl1, tlx3* | NA |
| dI6 | Glycinergic | Commissural Local (CoLo)<br>*- Satou et al. 2009, Kelly et al., 2023* | commissural Interneurons (cIN) | Secondary<br>*-Satou et al., 2009* | *lhx1, dmrt3, lbx1* | NA |
| V0v | Glutamatergic | Multipolar Commissural Descending (MCoD) and Unipolar Commissural Descending (UCoD)<br>*- Satou et al., 2012, Bernhardt et al., 1990, Kelly et al., 2023* | Dorsolateral commissural (dlc) and/or excitatory commissural Interneurons (ecINs) | Secondary<br>*-Satou et al., 2012* | ***evx1**, lhx1, nrn1,*<br>*developmental ***dbx1***<br>*- Juárez-Morales* | *cbln2+*<br>subcluster |
| V1 | Glycinergic/<br>GABAergic | Circumferential Ascending (CiA)<br>*- Higashijima et al., 2004, Kelly et al., 2023* | ascending Interneurons (aINs)<br>*- Li et al., 2004* | Secondary | ***en1,** lhx1, pnoc, pax2, sp9, foxd3, gbx2, otp* | NA |
| V2a | Glutamatergic/<br>Cholinergic | Circumferential Descending (CiD)<br>*- Kimura et al., 2006, Kelly et al., 2023, Kelly et al., 2023* | descending Interneurons (dINs)<br>*- Soffe et al., 2009* | Secondary<br>*- Bernhardt et al., 1990* | ***vsx2**, nrn1, nkx6-2, lhx3* | *chat- sox21+, prdm8+, sox14+*<br>subcluster |
| V2b | GABAergic | Ventral Longitudinal Descending (VeLD)<br>*- Kumura et al., 2006, Kelly et al., 2023* | descending Interneurons (dINs)<br>*- Soffe et al., 2009, Li et al., 2004* | Primary<br>*- Bernhardt et al., 1990* | *gata2, gata3, lhx1, sox1, tal1* | *slc6a5+*<br>subcluster |
| KA' | GABAergic | Dorsal Kolmer-Agduhr (KA')<br>*- Yang et al., 2020* | Kolmer-Agduhr Interneuron<br>*- Roberts and Clarke 1982, Kale et al., 1987, Binor and Heathcote, 2001* | Secondary | *pkd1l2, kctd8, sox1, tal1* | NA |
| KA" | GABAergic/<br>Dopaminergic | Ventral Kolmer-Agduhr (KA")<br>*- Yang et al., 2020* | Kolmer-Agduhr Interneuron<br>*- Roberts and Clarke 1982, Kale et al., 1987, Binor and Heathcote, 2001* | Secondary | *pkd1l2, sox1, nkx6-2, foxa2, tal1* | *chgb, cfap210, sall2, daw1, cdx4* |
| ? | Glycinergic | NA | NA | NA | *megf11+, lhx1* | NA |
| ? | Glycinergic | NA | NA | NA | *foxp2+, pax2+* | NA |
| Reviewed in: | | *Lewis and Eisen 2003, Sengupta and Bagnall, 2023, Wilson and Sweeney, 2023, and Cucun et al., 2024. Reviewed in mouse in Delile et al., 2019* | *Roberts, 2011* | *Saint-Amant and Drapeau 2001* | | |

**Table S1. Conserved markers from zebrafish can be used to identify *Xenopus* neuron counterparts based on literature analysis.** Cardinal neurons have been identified in zebrafish literature by their neurotransmitter use, morphological classification, and birth date. These identities can be tied to their likely *Xenopus* counterpart in our scRNA-seq dataset based on cluster Marker Genes. Bold genes indicate marker genes that have been previously identified in *Xenopus.* For some neurons, additional subclusters could be identified for future studies. Two clusters of glycinergic neurons remain unidentified, and V3 and dI1-5 INs could not be found in our dataset.

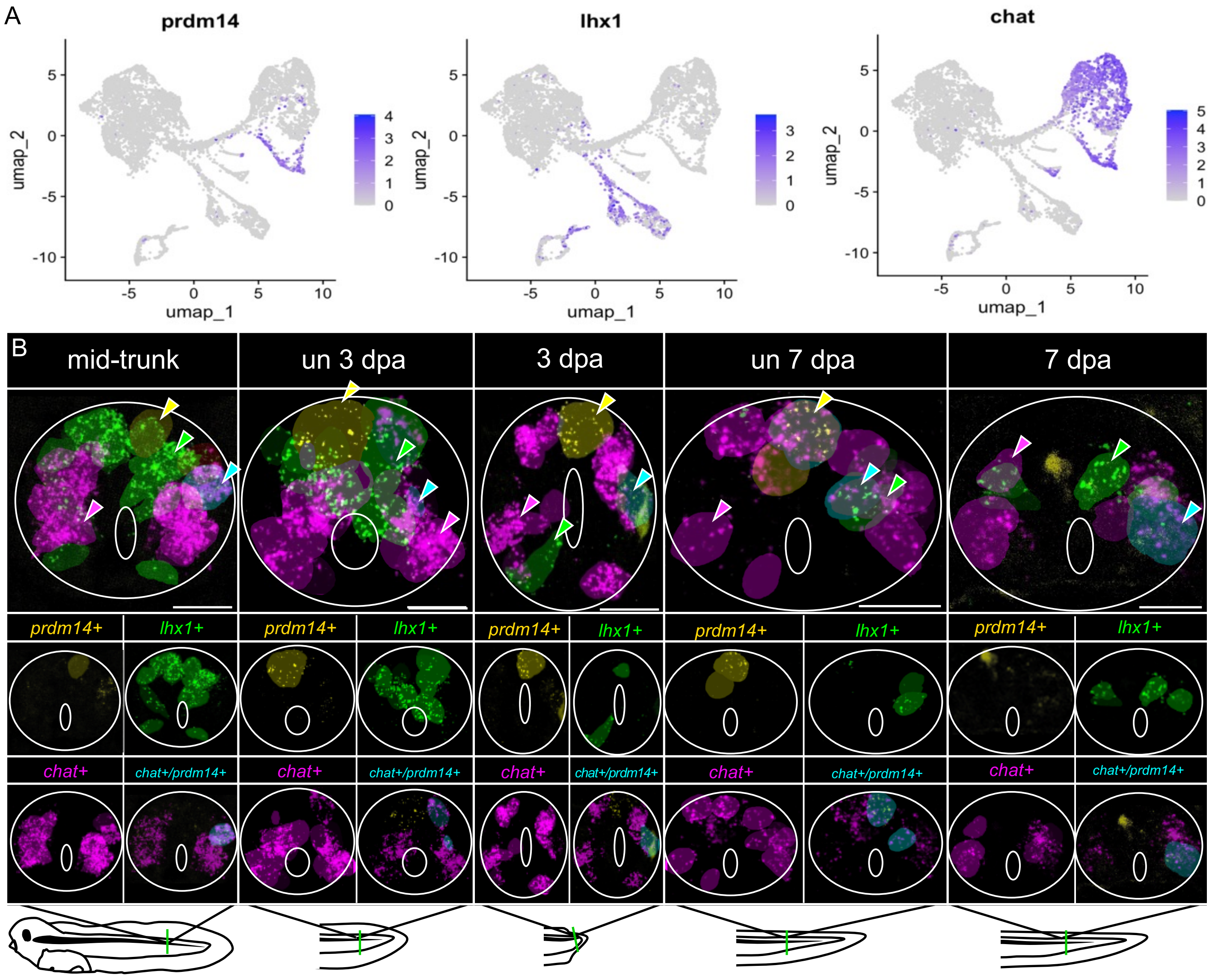


**Figure S3. HCR of neuron types within regenerating tissue.** A) UMAPs showing chosen HCR probe gene expression of *prdm14, lhx1,* and *chat*. B) Images showing the raw HCR Z-stacks used to generate masks in Figure 3. Segmented cell masks are overlayed. White solid line denotes spinal cord boundary (outer) and central canal (inner). HCR: yellow is *prdm14*, green is *lhx1*, magenta is *chat*. Cells: yellow is *prdm14* + RBNs, green is *lhx1*+ INs, magenta is *prdm14-/chat*+ MNs*,* cyan is *prdm14+/chat*+ MNs. Example cells of each type are shown with a color matched arrowhead. All scale bars are 100 μm.

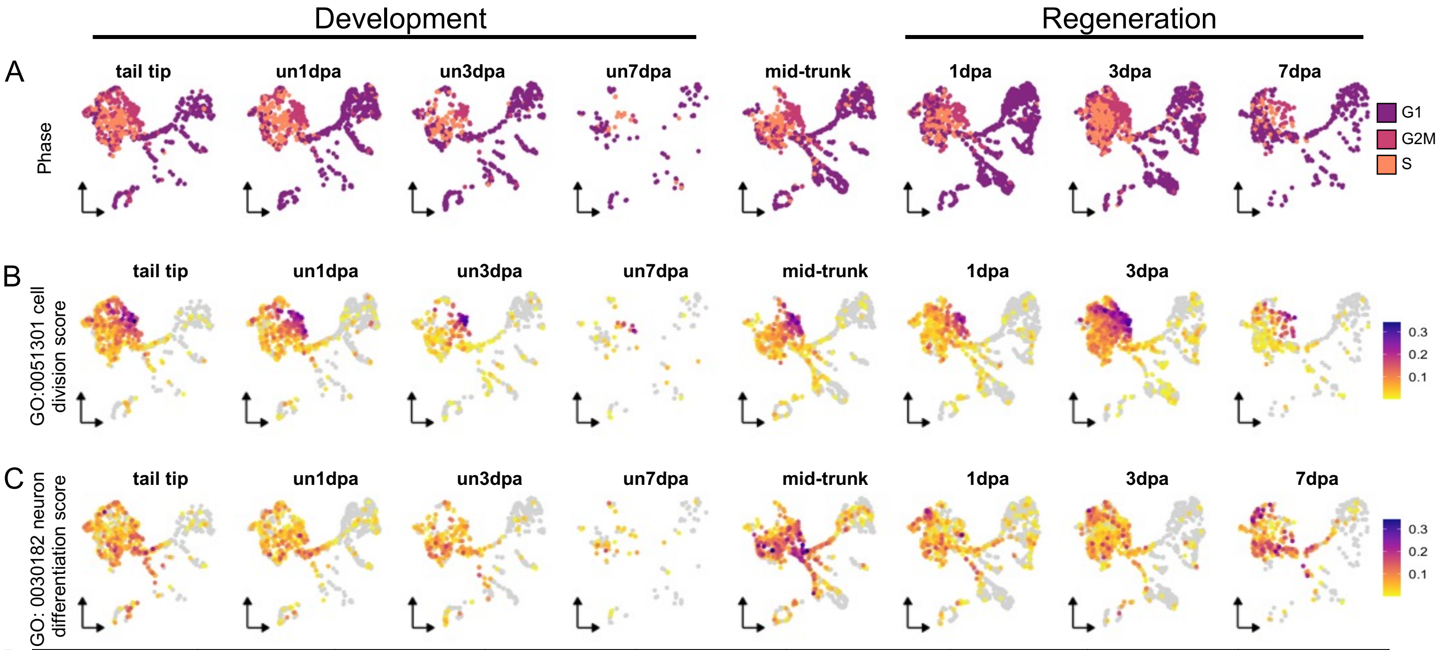


D

| | tail tip | un1dpa | un3dpa | un7dpa | mid-trunk | 1dpa | 3dpa | 7dpa |
|---|---|---|---|---|---|---|---|---|
| tail tip | | **** | **** | **** | **** | **** | *** | **** |
| un1dpa | **** | | ns | ns | ns | **** | **** | ns |
| un3dpa | **** | ns | | ns | *** | ns | **** | ns |
| un7dpa | **** | ns | ns | | ns | ns | **** | ns |
| mid-trunk | * | **** | **** | **** | | **** | **** | ** |
| 1dpa | **** | ns | ns | ns | **** | | **** | * |
| 3dpa | **** | ns | ns | ns | **** | ns | | **** |
| 7dpa | *** | **** | **** | **** | ns | **** | **** | |

neuron differentiation GO term →

cell division GO term

**Figure S4. Faceted UMAPs of cell processes throughout development and regeneration.** A) Cell cycle phase (G1, G2M, or S) over scRNA-seq timepoints. B) GO:0051301 cell division module score over scRNA-seq timepoints. C) Neuron Differentiation Module Score over scRNA-seq timepoints. D) Comparative adjusted p-values for neuron differentiation GO term (bottom left) and cell division GO term (top right) over developmental and regenerative time.

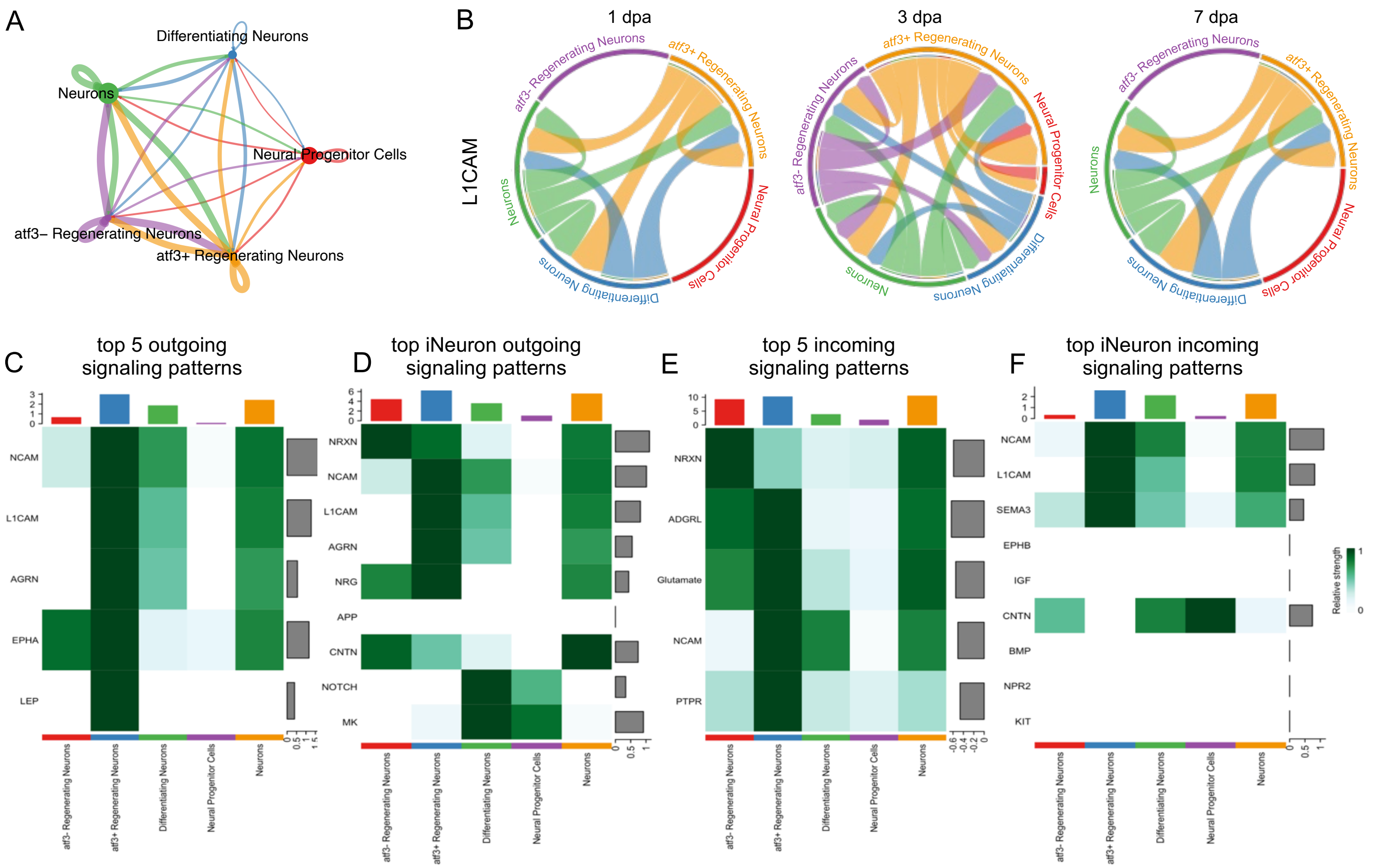


**Figure S5. *atf3*+ Regenerating Neurons do not represent a major signaling hub but do show similar top signaling pathways to zebrafish iNeurons.** A) Comparative signaling strength of regenerating neurons. B) L1CAM signaling during regeneration. C) Top 5 outgoing signaling patterns at 1 dpa D) Visualization of the top outgoing signaling patterns identified zebrafish iNeurons during regeneration in *Xenopus* regenerating neurons. E) Top 5 incoming signaling patterns at 1dpa F) Visualization of the top incoming signaling patterns identified zebrafish iNeurons during regeneration in *Xenopus* regenerating neurons.

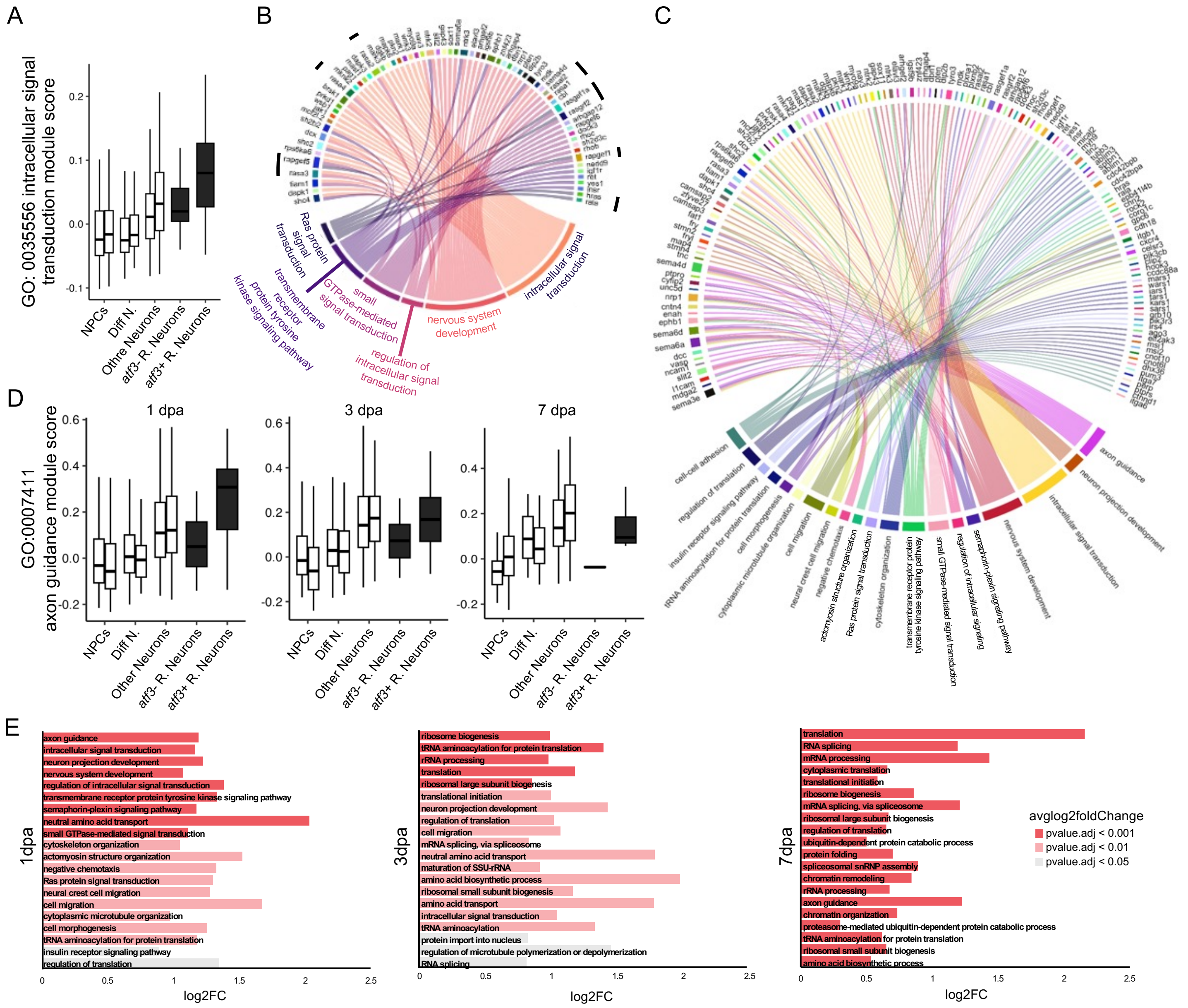


**Figure S6. Top 20 GO terms and associated genes in *atf3*+ Regenerating Neurons at 1 dpa compared to 3 and 7 dpa.** A) Plot of GO:0035556 intracellular transduction score at 1 dpa in different cell clusters in uninjured (left bar) or regenerating (right bar) timepoints. Black denotes regeneration-specific subclusters. B) Plot of GO:0035556 intracellular transduction score at 1dpaand associated modules at 1 dpa (bottom), compared to upregulated genes representing scores (top). Black bars indicate Ras associated proteins, which represent one of the top enriched gene families. C) Plot of all top 20 GO terms and associated genes at 1dpa. D) Plot of GO:0035556 intracellular transduction score in different cell clusters in uninjured (left bar) or regenerating (right bar) timepoints at 1 dpa (left), 3 dpa (middle), and 7 dpa (right). Black denotes regeneration-specific subclusters. E) Top 20 GO terms upregulated in *atf3*+ regenerating neurons compared to other Neurons at 1 dpa (left), 3 dpa (middle), and 7 dpa (right).

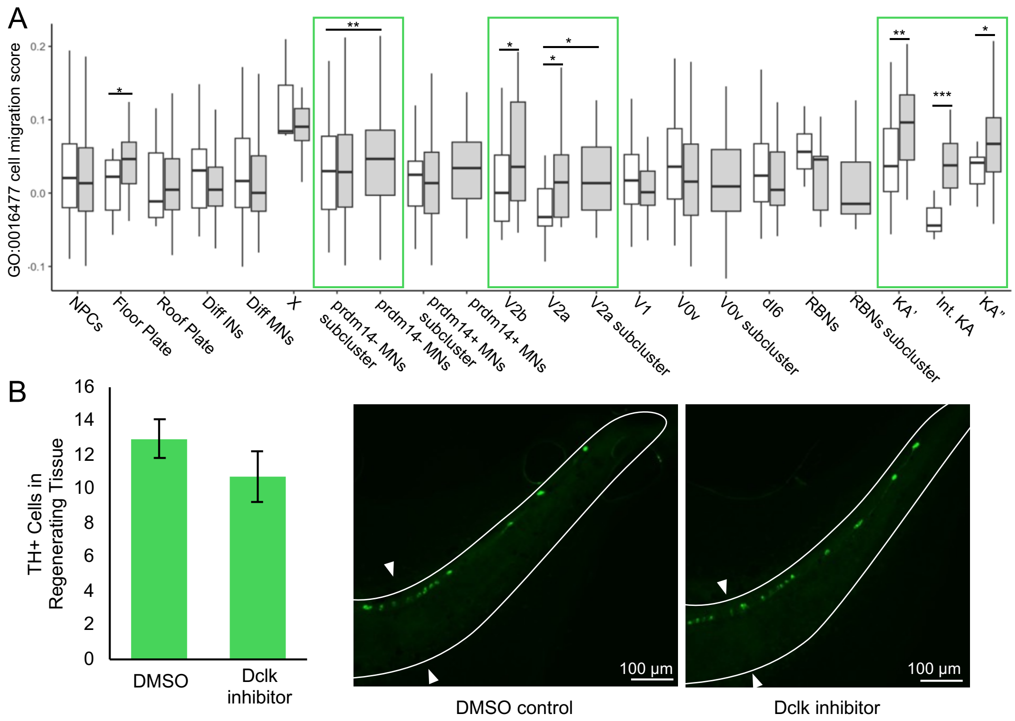


**Figure S7. Cell migration may underlie mature neuron displacement.** A) GO:0016477 cell migration scores are enriched in regenerating stages in regeneration-specific *prdm14-* MNs, V2bs, V2as, and KA INs (green rectangles). *p-value < 0.05; **p-value < 0.01; ***p-value < 0.001 by Wilcoxon test. B) Dclk inhibition slightly reduces the number of TH+ KA" INs in the regenerating tail (not significant via Wilcoxon test, n = 5 each).
